# Microtubules Promote the Non-cell Autonomy of MicroRNAs by Inhibiting their Cytoplasmic Loading into ARGONAUTE1 in *Arabidopsis*

**DOI:** 10.1101/2021.05.26.445899

**Authors:** Lusheng Fan, Cui Zhang, Yong Zhang, Ethan Stewart, Jakub Jez, Keiji Nakajima, Xuemei Chen

## Abstract

Mobile microRNAs (miRNAs) serve as local and long-distance signals in developmental patterning and stress responses in plants. However, mechanisms governing the non-cell autonomous activities of miRNAs remain elusive. Here, we show that mutations that disrupt microtubule dynamics are specifically defective for the non-cell autonomous actions of mobile miRNAs, including miR165/6 that is produced in the endodermis and moves to the vasculature to pattern xylem cell fates in *Arabidopsis* roots. We show that KTN1, a subunit of a microtubule-severing enzyme, is required in source and intermediary cells to inhibit the loading of miR165/6 into ARGONUATE1 (AGO1), which is cell-autonomous, to enable the miRNA‟s cell exit. Microtubule disruption enhances the association of miR165/6 with AGO1 in the cytosol. These findings suggest that, while cell-autonomous miRNAs load into AGO1 in the nucleus, cytoplasmic AGO1 loading of mobile miRNAs is a key step regulated by microtubules to promote the range of miRNA‟s cell-to-cell movement.

## INTRODUCTION

MicroRNAs (miRNAs) are 20-24-nucleotide (nt) long, non-coding RNAs that regulate nearly all aspects of plant life via posttranscriptional gene silencing (Bologna and Voinnet, 2014; Yu et al., 2019). In plants, the entire miRNA biogenesis process including *MIR* gene transcription, primary miRNA (pri-miRNA) processing, methylation of miRNA/miRNA* duplexes and their loading into ARGONUATE1 (AGO1), and the formation of RNA-induced silencing complexes (RISCs) is thought to occur in the nucleus. miRISCs are then exported to the cytoplasm (Bologna et al., 2018), where they recognize target mRNAs via a high degree of miRNA-mRNA sequence complementarity, leading to (1) transcript cleavage through the endonuclease activity of AGO1, or (2) translational inhibition*(Baumberger and Baulcombe, 2005; Brodersen et al., 2008; Li et al., 2013)*.

A key feature of RNA silencing mediated by small RNAs is non-cell autonomy, in which a silencing signal moves locally (cell to cell) and systemically (over a long distance) (Chitwood et al., 2009; Klesen et al., 2020; Melnyk et al., 2011). Small RNAs, including transgene and endogenous small interfering RNAs (siRNAs) and miRNAs, have been recognized as the mobile signals (Buhtz et al., 2010; Lewsey et al., 2016; Li et al., 2021; Lin et al., 2008; Molnar et al., 2010; Zhang et al., 2014). Mobile small RNAs may coordinate events in distant plant parts. For example, several miRNAs are induced in shoots and transported to roots to regulate responses to nutrient starvation and nodulation (Buhtz et al., 2010; Lin et al., 2008; Okuma et al., 2020; Pant et al., 2008). Several mobile small RNAs, such as miR165/6, miR394, and the trans-acting siRNA tasiR-ARF, serve as morphogenic signals that pattern cell fates in development (Carlsbecker et al., 2010; Chitwood et al., 2009; Knauer et al., 2013; Miyashima et al., 2011). In roots, *MIR165/6* genes are specifically expressed in the endodermis, but miR165/6 forms a gradient towards the center of the vasculature, which leads to an opposing gradient in the expression of its target gene *PHABULOSA* (*PHB*) (Carlsbecker et al., 2010; Miyashima et al., 2011). Cells in the vasculature with the highest level of *PHB* expression become metaxylem while neighboring cells with lower levels of *PHB* expression take on the protoxylem fate. The mobile agents in non-cell autonomous RNA silencing by transgenes are likely siRNA duplexes (Devers et al., 2020). As AGO1 is cell-autonomous, the movement of siRNA duplexes from cell to cell is negated by their loading into AGO1 during transit (Devers et al., 2020). Previous studies show that miRNAs spread through the plasmodesmata in *Arabidopsis* and generate a gradient distribution pattern consistent with passive diffusion between cells (Chitwood et al., 2009; Skopelitis et al., 2017; Vaten et al., 2011). However, it was also observed that the cell-to-cell movement of miRNAs is directional at some cell-cell interfaces, implicating regulated mobility (Skopelitis et al., 2018). A modeling approach showed that diffusion alone cannot account for the miR165/6 activity gradient (Muraro et al., 2014). Mechanisms that regulate the cell-to-cell movement of small RNAs are largely unknown.

Microtules (MTs) are dynamic structures with a shrinking minus end and a growth-biased, dynamic and unstable plus end, which together contribute to MT displacement and reorientation, also known as treadmilling. MT dynamics is tightly regulated by MT-associated proteins (MAP), such as MOR1, which is required for the rapid shrinkage and growth of MTs (Kawamura and Wasteneys, 2008). The *KATANIN1* (*KTN1*) gene of *Arabidopsis* encodes the p60 subunit of *Arabidopsis* Katanin, which severs MTs at crossover sites to maintain organized MT arrays and to ensure dynamic MT re-organization in the cytosol (Lin et al., 2013). The dynamic MT cytoskeleton plays pivotal roles in multiple fundamental cellular processes (Ehrhardt and Shaw, 2006), including the cell-to-cell movement of *Tobacco Mosaic Virus* (Boyko et al., 2007). Interestingly, *KTN1* was also found to be required for the translation repression, but not the RNA cleavage, activity of plant miRNAs, suggesting that MT dynamics regulates miRNA activities (Brodersen et al., 2008). It is unknown whether MTs are required for the cell-to-cell movement of miRNAs.

Here, we show that mutations in *KTN1* and *MOR1* compromise the cell-to-cell movement miRNAs in leaves and roots. In roots, the mutants exhibit xylem patterning defects reminiscent of reduced mobility of miR165/6, which is also reflected by a shallower *PHB* gradient. More importantly, we show that, while critical for the non-cell autonomous action of miR165/6, *KTN1* is dispensable for the cell-autonomous activity of the miRNA. Furthermore, *KTN1* is required in cells that miR165/6 is made in (endodermis) or transit through (protoxylem), but dispensable in destination cells (metaxylem). Through tissue-specific analysis of AGO1 loading, we found that *KTN1* inhibits the loading of miR165/6 into AGO1 in the source tissue, thus promoting its exit. Disruption of MTs enhances the loading of miR165/6 into AGO1 in the cytoplasm. These findings reveal that, while cell-autonomous miRNAs undergo AGO1 loading in the nucleus, cytoplasmic AGO1 loading is a key point of regulation for mobile miRNAs and that MTs inhibit cytoplasmic RISC formation to promote the non-cell autonomy of miRNAs.

## RESULTS

### Identification of KTN1 as a Potential Regulator of the Non-cell Autonomous Activities of miRNAs

We performed an ethyl methanesulfonate mutagenesis screen using the *SUC2::amiR-SUL* transgenic line, in which the amiR-SUL artificial miRNA is specifically produced in companion cells in the phloem and non-cell autonomously silences the *SULFUR* (*SUL*) gene in leaf mesophyll cells, leading to vein-centered leaf chlorosis (de Felippes et al., 2011). We isolated a mutant with reduced leaf bleaching, which implies compromised amiR-SUL activity (Figure 1A). The mutant exhibited morphological phenotypes reminiscent of mutants in *KATANIN1* (*KTN1*), such as smaller and rounder leaves (Figure 1A). Indeed, a point mutation at the splice site of the first intron of *KTN1* was identified in this mutant and termed *ktn1-20* (Figure S1A). Another *ktn1* mutant with a T-DNA insertion, *ktn1-2* (Figure S1A), also led to weaker chlorosis when introduced into *SUC2::amiR-SUL* (Figure 1A). A genomic fragment containing the promoter and coding region of *KTN1* fully rescued the developmental and leaf chlorosis phenotypes of *SUC2::amiR-SUL ktn1-20*, further confirming that loss of function in *KTN1* was responsible for these phenotypes (Figure 1B).

**Figure 1.**
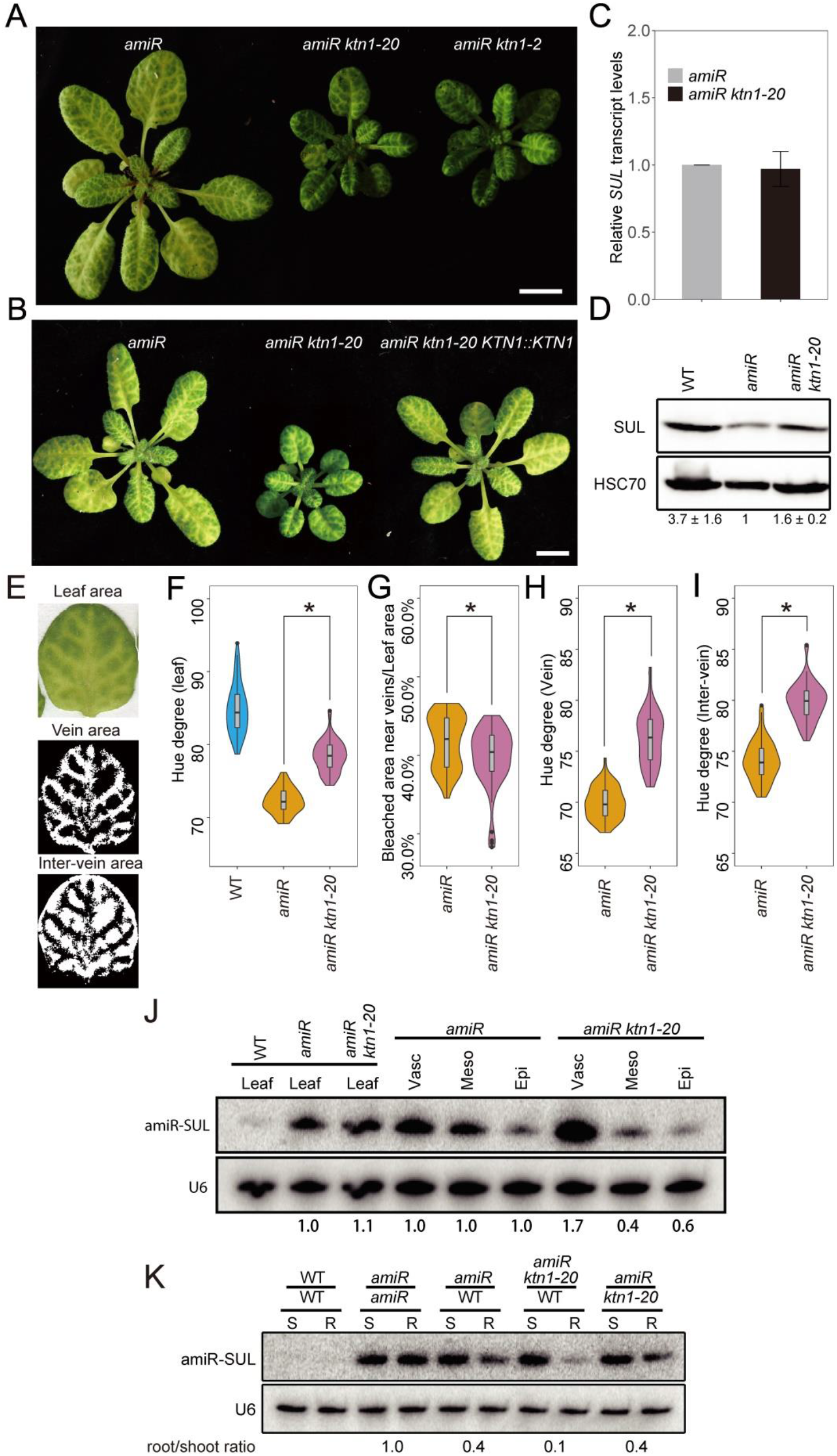
*KTN1* Is Required for Intercellular and Systemic Movement of amiR-SUL. (A) 4-week-old plants of *SUC2::amiR-SUL* (*amiR*)*, amiR ktn1-20* and *amiR ktn1-2.* (B) 4-week-old plants of *amiR, amiR ktn1-20 and amiR ktn1-20 KTN1::KTN1,* showing that *KTN1::KTN1* fully rescues the developmental and leaf bleaching phenotypes of *ktn1-20.* Scale bars in (A) and (B), 1 cm. (C) Relative levels of the *SUL* transcript in *amiR* and *amiR ktn1-20.* Error bars represent SD calculated from three biological replicates, each with three technical replicates. *UBQ5* RNA served as the internal control. (D) SUL protein in *amiR* and *amiR ktn1-20* as determined by protein gel blot analysis with anti-SUL antibodies. HSC70 served as the loading control. Relative protein levels are indicated below the gel images by numbers representing mean+/−SD calculated from three biological replicates. (E) Representative images used to quantify areas of leaf chlorosis. (F) Hue color value representing the levels of chlorosis for whole leaves in wild type (WT), *amiR* and *amiR ktn1-20.* Lower values correspond to increased chlorosis. 60 degrees represent yellow and 120 degrees represent green. (G) Quantification of the area of vein-centered bleaching normalized against total leaf area in *amiR* and *amiR ktn1-20.* (H and I) Quantification of levels of chlorosis (indicated by hue value) for vein and inter-vein regions in *amiR* and *amiR ktn1-20. p*-values were calculated by Student‟s *t* test. *, *p-*value < 0.01. 60 leaves from 10 individual plants were analyzed for *amiR* and *amiR ktn1-20* in (F) to (I). (J) RNA gel blot analysis of amiR-SUL in Meselect-separated vascular, mesophyll and epidermal tissues. U6 was used a loading control. The amiR-SUL levels in *SUC2::amiR-SUL ktn1-20* were relative to those in *SUC2::amiR-SUL* in each tissue and indicated by the numbers below the gel images. (K) RNA gel blot analysis of amiR-SUL in scions (S) and rootstocks (R) of WT, *amiR*, *amiR ktn1-20*, *and ktn1-20* in the indicated grafting combinations. The values below the gel images represent root/shoot ratios of amiR-SUL relative to that of *amiR*/amiR set as 1.0. See also Figure S1 and S2.

The weaker leaf chlorosis of *SUC2::amiR-SUL ktn1-20* could be due to compromised miRNA biogenesis. RNA gel blot assays showed that the levels of amiR-SUL and miR168 were similar between *SUC2::amiR-SUL ktn1-20* and *SUC2::amiR-SUL* (Figure S1B). Small RNA-seq with *ktn1-20* and wild type did not find global changes in miRNA abundance in the mutant (Figure S1 C-E). These results ruled out a role of *KTN1* in miRNA biogenesis. Real-time RT-PCR analysis showed that *SUL* transcript levels were similar in *SUC2::amiR-SUL ktn1-20* and *SUC2::amiR-SUL* (Figure 1C). The levels of the SUL protein were lower in *SUC2::amiR-SUL* than in wild type (Col-0), as expected, and higher in *SUC2::amiR-SUL ktn1-20* than in *SUC2::amiR-SUL* (Figure 1D). Thus, these data are consistent with previous findings that *KTN1* mediates the translation repression activity of plant miRNAs (Brodersen et al., 2008).

We next asked whether the *ktn1-20* mutation resulted in a reduced range of movement of amiR-SUL, which would be reflected by a narrow span of bleaching around the veins. We used Open CV (Bradski and Kaehler, 2000) to quantify the spatial patterns of leaf bleaching (Figure 1E; see Methods). At the level of whole leaves, the severity of bleaching was significantly alleviated in *SUC2::amiR-SUL ktn1-20* relative to *SUC2::amiR-SUL*, consistent with visual inspection (Figure 1F). The area of bleaching along the vasculature, representing the range of non-cell autonomous action of amiR-SUL, was significantly reduced in the *SUC2::amiR-SUL ktn1-20* mutant, even after normalization to total leaf area (Figure 1G). Interestingly, in addition to reduced bleaching near the veins, the level of bleaching in inter-vein areas was also remarkably decreased in *SUC2::amiR-SUL ktn1-20* (Figure 1H and 1I). This suggests that amiR-SUL‟s range of movement in *SUC2::amiR-SUL* was likely broader than the distance of 15 cells from the veins as previously reported (de Felippes et al., 2011). The strong silencing near veins and weaker silencing in inter-vein regions in *SUC2::amiR-SUL* are reminiscent of the threshold effects observed for the activities of mobile miR165/6 and tasiR-ARF in leaves (Chitwood et al., 2009; Merelo et al., 2016; Skopelitis et al., 2017).

To definitively show that *KTN1* is required for the cell-to-cell movement of amiR-SUL, we mechanically separated leaf vascular, mesophyll and epidermal tissues by Meselect (Svozil et al., 2016) and examined amiR-SUL levels in source (vascular) and recipient (mesophyll and epidermal cells) tissues. Tissue enrichment was successful, as shown by the expression of *SUC2*, *CAB3*, and *ATML1*, which mark vascular, mesophyll, and epidermal tissues, respectively (Figure S2). As expected, amiR-SUL was enriched in the vascular tissue in both *SUC2::amiR-SUL* and *SUC2::amiR-SUL ktn1-20* as compared to mesophyll and epidermal tissues. More importantly, the abundance of amiR-SUL was increased in the vascular tissue but decreased in mesophyll and epidermal tissues in *SUC2::amiR-SUL ktn1-20* relative to *SUC2::amiR-SUL* (Figure 1J), indicating reduced cell-to-cell movement of amiR-SUL in the *ktn1-20* background.

We employed *Arabidopsis* micrografting to examine whether *KTN1* impacts the long-distance movement of amiR-SUL. In wild type rootstocks micrografted onto *SUC2::amiR-SUL* scions (*SUC2::amiR-SUL*/WT), a substantial amount of amiR-SUL was detected, suggesting that amiR-SUL was able to move from shoot to root. The levels of amiR-SUL were similar in *SUC2::amiR-SUL*/WT vs. *SUC2::amiR-SUL*/*ktn1-20* rootstocks, suggesting that the long-distance trafficking of amiR-SUL was not impacted when the *ktn1-20* mutation was in the recipient tissue. Interestingly, the levels of amiR-SUL in *SUC2::amiR-SUL ktn1-20*/WT rootstocks was substantially lower than those in *SUC2::amiR-SUL*/WT rootstocks (Figure 1K). These results demonstrated that *KTN1* is required in incipient tissues for the long-distance movement of amiR-SUL.

### KTN1 Patterns Xylem Cell Fates by Enabling the Non-cell Autonomous Function of miR165/6

We next evaluated whether *KTN1* promotes the non-cell autonomous activities of endogenous miRNAs. The well-characterized non-cell autonomy of miR165/6 and the dosage-dependent activity of the miRNA in the regulation of its target gene *PHB* leading to xylem patterning (Carlsbecker et al., 2010; Miyashima et al., 2011) provided a superb model for our purposes. First, we examined the xylem patterns in *ktn1-20*. There are two metaxylem files flanked by two protoxylem files in a wild type root (Figure 2A). However, in *ktn1-20* roots, metaxylem was present, but protoxylem differentiation was severely impaired (Figure 2A and 2B). In *ktn1-20* roots, protoxylem was replaced by metaxylem (Figure 2A), or ectopic files of metaxylem were found (Figure 2B). Expression of *KTN1* under its native promoter fully rescued the xylem patterning defects, indicating that the impaired xylem development was due to the *ktn1-20* mutation (Figure 2A and 2B). As xylem differentiation is controlled by *PHB* in a dosage-dependent manner: high and low levels of *PHB* specify metaxylem and protoxylem, respectively (Carlsbecker et al., 2010; Miyashima et al., 2011), we compared the *PHB* expression patterns between WT and *ktn1-20*. In WT, PHB-GFP distribution formed a gradient across the stele, with the highest level in the center and a drastic decline towards the periphery (Figure 2C and 2E). In contrast, the sharp gradient distribution was disrupted in *ktn1-20*, with more cell files showing high levels of PHB-GFP (Figure 2D and 2E), which was consistent with the higher numbers of metaxylem. To rule out that miR165/6 biogenesis was affected in *ktn1-20*, we first examined the spatial expression patterns of *MIR165A*, *MIR165B*, *MIR166A* and *MIR166B*, the four out of nine *MIR165*/*6* genes with detectable expression in the root (Miyashima et al., 2011). The promoters of these genes drove the expression of GFP specifically in the endodermis in the root (Figure S3A). The *ktn1-20* mutation did not affect the tissue-specific promoter activities of the four genes (Figure S3A and S3B). The levels of all nine pri-miR165/6s were determined by qRT-PCR, and no significant changes were found for any transcript between WT and *ktn1-20* (Figure. S3C). RNA gel blot assays showed that miR165/6 accumulation was similar between WT and *ktn1-20* (Figure S3D), which was consistent with results from small RNA sequencing (Figure S1F). Taken together, these results indicate that *KTN1* is required for the non-cell autonomous function of miR165/6, but dispensable for its biogenesis.

**Figure 2.**
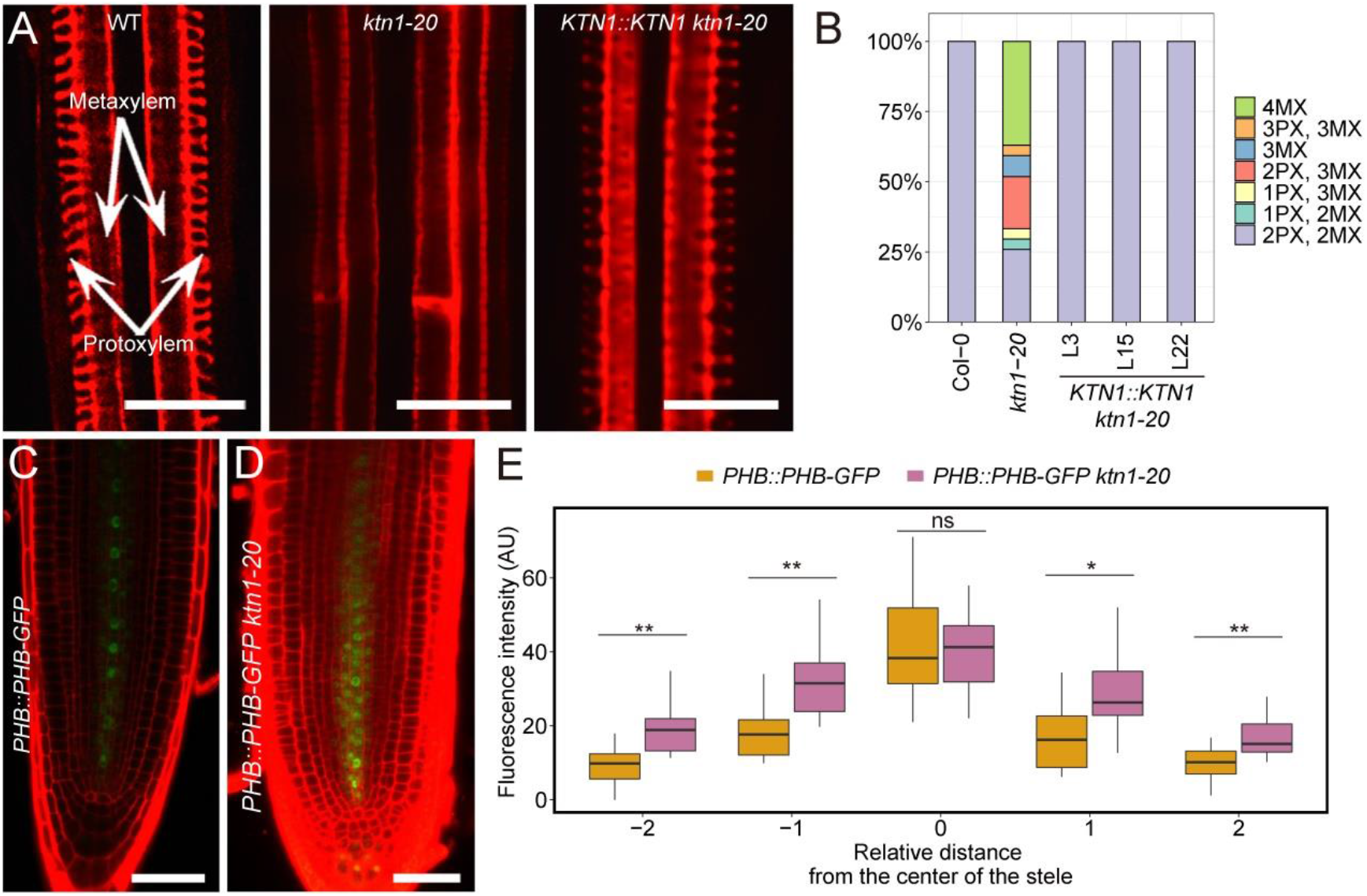
Defects in Xylem Patterning and PHB-GFP Gradient Distribution in the *ktn1-20* Mutant. (A) Xylem patterns in WT, *ktn1-20* and the complementation line *KTN1::KTN1 ktn1-20*. The white arrows indicate the two metaxylem layers flanked by two protoxylem layers. In the *ktn1-20* image, four layers of metaxylem are seen. (B) Quantification of xylem phenotypes in WT (Col-0), *ktn1-20* and *KTN1::KTN1 ktn1-20*. 20-30 individual roots were analyzed. In some roots of the *ktn1-20* mutant, xylem cell wall morphology was intermediate between protoxylem and metaxylem such that the spiral cell wall thickening pattern was not as striking. Such xylem was counted as metaxylem, as an obvious cell-cell boundary was found between two consecutive cells, while such boundary is not present in protoxylem. (C, D) Expression patterns of *PHB::PHB-GFP* in WT and *ktn1-20*. Scale bars, 20 μm in (A); 50 μm in (C) and (D). (E) Quantification of PHB-GFP signal intensity measured across the root diameter in WT and *ktn1-20*. 15-20 individual roots were imaged for quantification. AU: arbitrary unit. *p*-values were calculated by Student’s *t* test. *\*, p-*value < 0.05, **; *p-*value < 0.01; ns, not significant. See also Figure S3, S4 and S5.

### MOR1 Is Required for the Non-cell Autonomous Actions of miRNAs

As *KTN1* is required for the formation of highly ordered microtubule arrays, we further examined the effects of another MT mutant, *mor1-1*, on the non-cell autonomous activities of miRNAs. The *mor1-1* allele is temperature sensitive - it fails to form intact MTs only at the restrictive temperature (30℃) (Whittington et al., 2001). *SUC2::amiR-SUL mor1-1* showed no obvious leaf bleaching defects when grown under normal temperature (Figure S4A). However, when the plants were transferred to the restrictive temperature, newly emerged leaves showed reduced leaf bleaching in comparison to *SUC2::amiR-SUL* (Figure S4B), while amiR-SUL levels in *SUC2::amiR-SUL mor1-1* showed no significant changes relative to *SUC2::amiR-SUL* under either normal or restrictive temperature (Figure S4C).

We further examined the xylem patterns of *mor1-1* grown under the restrictive temperature. Xylem differentiation into metaxylem and protoxylem was not affected in WT at 30°C(Figure S5, A and S5B). In contrast, metaxylem formed in the position of protoxylem or was increased in number in *mor1-1* under the restrictive temperature (Figure S5A and S5B). The PHB-GFP gradient in the stele was properly established in WT under the restrictive temperature (Figure S5C and S5D). However, the graded distribution of PHB-GFP was abolished in *mor1-1* under the restrictive temperature, with more cell files expressing high levels of PHB-GFP (Figure S5C and S5D), consistent with the increased number of metaxylem cell files. The levels of miR165/6 were similar between WT and *mor1-1* under both normal and restrictive conditions (Figure S5E).

It has been shown that *ktn1* mutant roots are wider than wild type roots (Webb et al., 2002), which was also observed for *ktn1-20* roots (Figure S5F and S5G). We considered the possibility that the enlarged root diameter rather than the disruption of MTs *per se* prevented miR165/6 movement into the center of the stele. The temperature-sensitive nature of *mor1-1* provided an opportunity to examine the PHB-GFP gradient without the complication of enlarged root diameter. The graded PHB-GFP distribution was already disrupted in *mor1-1* (Figure S5C and S5D) at 16 hours after plants were transferred to the restrictive temperature when no change in root diameter was observed (Figure S5H and S5I). Note that the MT defects of *ktn1* and *mor1* mutants are not identical. In *ktn1* mutants, MTs are highly disorganized and show a network-like microtubule distribution (Lin et al., 2013), while in *mor1* mutants, MTs are fragmented (Whittington et al., 2001). Taken together, the effects of *ktn1* and *mor1* mutations on amiR-SUL and miR165/6 suggest that well-organized MTs or dynamic MTs play a vital role in the non-cell autonomous actions of miRNAs.

### KTN1 Is Dispensable for the Cell Autonomous Activity of miR165/6

Although *KTN1* was shown to enable the non-cell autonomous activity of miR165/6 in repressing its target *PHB* in the stele, *KTN1* does not necessarily promote the cell-to-cell movement of miR165/6. An alternative possibility is that *KTN1* is required for the target repression activity of miR165/6 in stele cells that receive this mobile miRNA. To test this possibility, we expressed *MIR165A* under the *CRE1* promoter, which is active specifically in the stele (Carlsbecker et al., 2010) where the miR165/6 target *PHB* is expressed. In the WT root, expression of *MIR165A* in the stele led to the conversion of metaxylem to protoxylem (Figure 3A), consistent with the notion that low level of *PHB* expression specifies protoxylem (Carlsbecker et al., 2010; Miyashima et al., 2011). As in WT, expression of *MIR165A* in the stele resulted in ectopic protoxylem formation in *ktn1-20* (Figure 3B). We further introduced *PHB::PHB-GFP* into both *CRE1::MIR165A* and *CRE1::MIR165A ktn1-20* and found that the graded distribution of PHB-GFP was disrupted in both wild type and *ktn1-20* backgrounds, with very weak PHB-GFP signals evenly distributed in the stele (Figure 3C–3E). These results demonstrated that miR165 was functional in *ktn1-20* when expressed in the same tissues as its target *PHB*. Therefore, *KTN1* is not required for the cell-autonomous activity of miR165/6. *KTN1*-regulated microtubule organization was reported to be required for the spiral cell wall thickening in the protoxylem (Schneider et al., 2021). Our results here showed that expression of *MIR165A* in the stele results in ectopic formation of protoxylem cell wall patterns in the *ktn1-20* background, which suggests that KTN1‟s role in protoxylem cell wall patterning is likely indirect, properly through its regulation of xylem cell fates via miR165/6.

**Figure 3.**
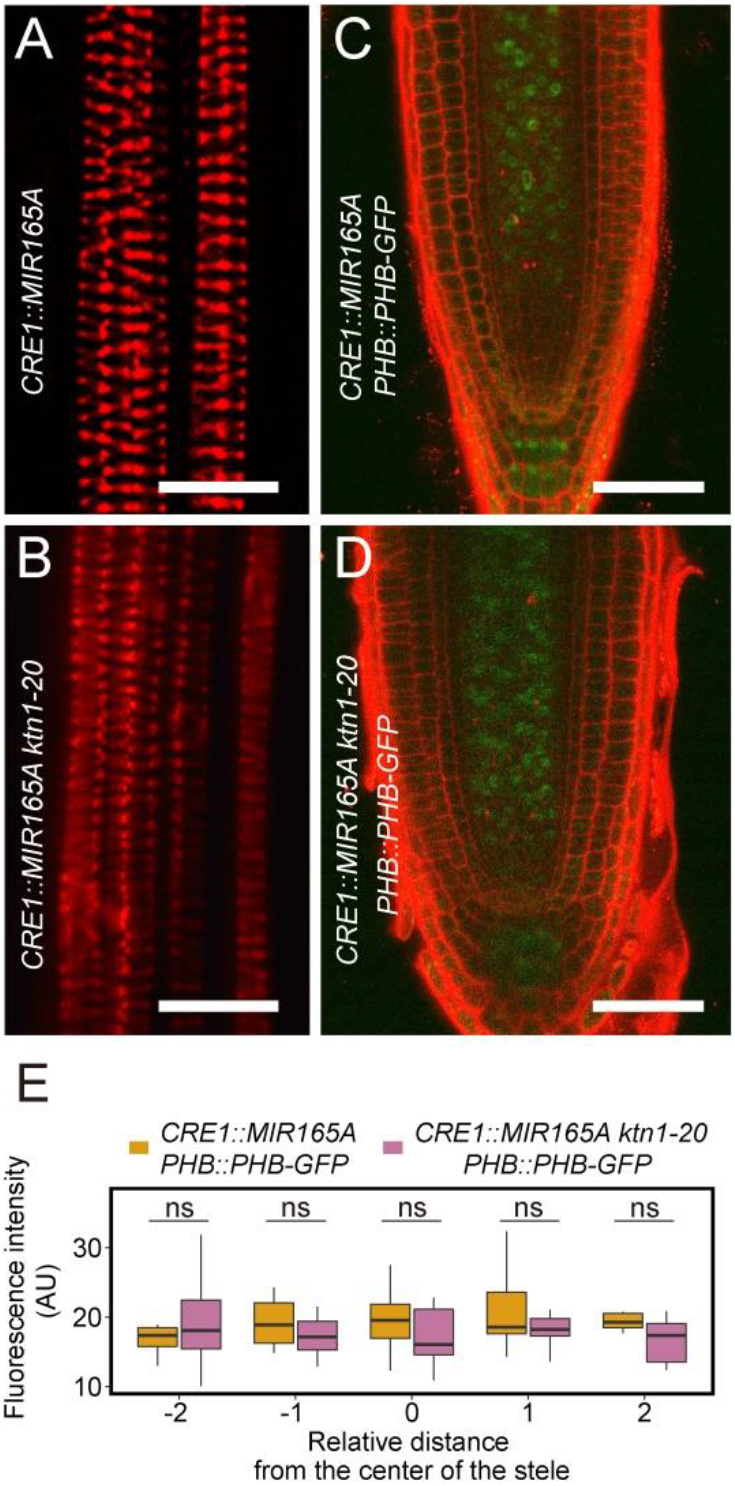
*KTN1* is dispensable for the cell-autonomous activity of miR165/6. (A, B) Xylem patterns in *CRE1::MIR165A* (A) and *CRE1::MIR165A ktn1-20* (B). Metaxylem-to-protoxylem transformation is seen in both genotypes. (C, D) Expression patterns of *PHB-GFP* in *pCRE1::miR165A* (C) and *pCRE1::miR165A ktn1-20* (D). Scale bars, 20 μm in (A) and (B); 50 μm in (C) and (D). (E) Quantification of PHB-GFP signal intensity across the root diameter in *pCRE1::miR165A* and *pCRE1::miR165A ktn1-20.* 10 individual roots were imaged for the quantification for each genotype. AU: arbitrary unit. ns, not significant.

### KTN1 Is Required in the Endodermis for the Non-cell Autonomous Action of miR165/6

To explore how *KTN1* promotes the non-cell autonomous action of miR165/6, we expressed *KTN1* in various root cell layers using layer-specific promoters, including *EN7* (endodermis), *AHP6* (protoxylem and adjacent pericycle), *ACL5* (metaxylem and procambia) and *SHR* (stele) (Figure 4A). Three independent T3 lines were analyzed for each construct. Among these transgenes, only *EN7::KTN1* fully rescued the xylem defects in *ktn1-20* as did *KTN1::KTN1* (Figure 4B and 4C). *AHP6::KTN1* only partially rescued the xylem patterning defects (Figure 4B and 4C). In addition, neither *ACL5::KTN1* nor *SHR::KTN1* was able to rescue the xylem defects in *ktn1-20* (Figure 4B and 4C). These results support the conclusion that *KTN1* is specifically required for the non-cell autonomous action of miR165/6 in the source tissue and in the tissue through which the miRNA transits.

**Figure 4.**
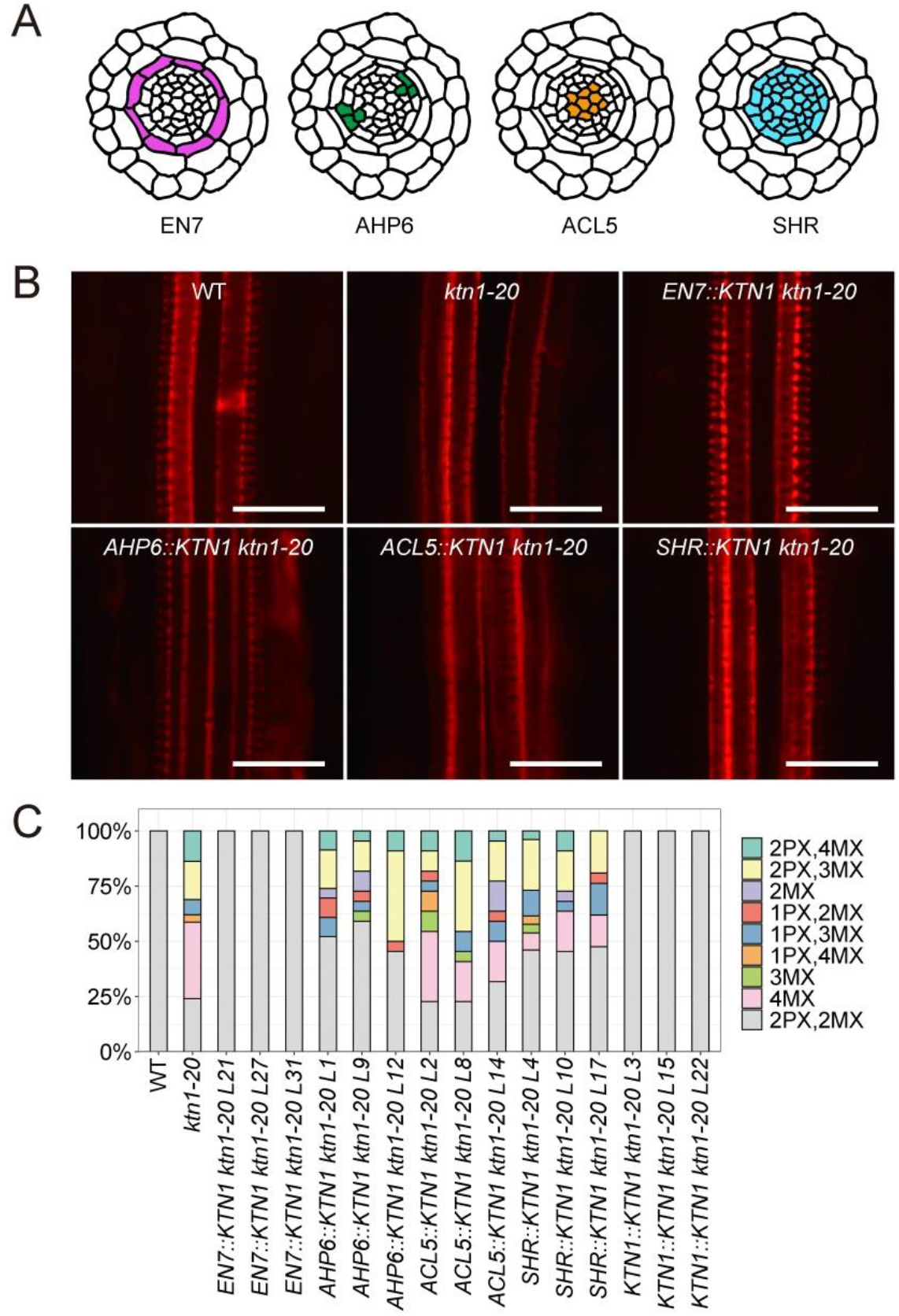
Expression of *KTN1* in the endodermis is crucial for xylem pattern formation. (A) Schematic representation of cross sections of an *Arabidopsis* root. Cell layers relevant to the tissue-specific expression of *KTN1* are colored. (B) Representative images showing xylem patterns in the indicated genotypes. Scale bars, 20 μm. (C) Quantification of the xylem phenotypes in the indicated genotypes. Three independent transgenic lines were analyzed for each transgene. 20-30 individual roots were included for each genotype.

### KTN1 Inhibits the Loading of miR165/6 into AGO1 in Source Cells to Promote its Cell-to-cell Movement

Previous studies show or implicate siRNA duplexes as the mobile agents in cell-to-cell movement (Devers et al., 2020). Furthermore, AGO1 expressed under various root layer-specific promoters are cell-autonomous (Brosnan et al., 2019); this study - see below), indicating that siRISCs or miRISCs are not cell-to-cell mobile. Thus, RISC formation is likely to inhibit the trafficking of small RNAs into neighboring cells. In fact, mobile siRNAs are increasingly depleted as they load into AGO1 during their cell-to-cell transit (Devers et al., 2020). Given this knowledge, we sought to determine whether KTN1 affects the loading of miR165/6 into AGO1 in source and recipient cells. We first generated transgenic lines expressing GFP-AGO1 under either *EN7* or *ACL5* promoters that are active in endodermis (Heidstra et al., 2004) and metaxylem/procambium (Muniz et al., 2008), respectively (Figure 5A). GFP-AGO1 signals were indeed restricted to the endodermis or metaxylem (Figure 5B), consistent with AGO1 being cell-autonomous. Immunoprecipitation (IP) was performed using an anti-GFP antibody followed by protein gel blots to confirm the successful IP of GFP-AGO1 (Figure S6A). RNAs were isolated from the immunoprecipitates and subjected to small RNA-seq. As compared to *Arabidopsis* total small RNA profiles showing a prominent 24-nt peak and a smaller 21-nt peak (Li et al., 2016), in GFP-AGO1 IP small RNA-seq, 21-nt small RNAs were enriched while 24-nt small RNAs were diminished (Figure S6B), indicating successful IP. The two biological replicates were highly correlated with each other for all the genotypes analyzed (Figure S6C-S6F). 10 differentially AGO1-associated miRNA species between *EN7::GFP-AGO1 ktn1-20* and *EN7::GFP-AGO1* (Figure S6G) and 19 differentially AGO1-associated miRNA species between *ACL5::GFP-AGO1 ktn1-20* and *ACL5::GFP-AGO1* (Figure S6H) were found. Among these miRNA species, AGO1-associated miR165/6 was at a higher level in the endodermis as compared to the metaxylem while AGO1-associated miR156 was at a lower level in the endodermis as compared to the metaxylem in both *ktn1-20* and wild type (Figure 5C), which is consistent with findings from a previous study (Brosnan et al., 2019). Strikingly, the abundance of AGO1-bound miR165/6 was higher in the endodermis, but lower in the metaxylem, in *ktn1-20* as compare to wild type (Figure 5C). These results suggest that *KTN1* suppresses the loading of miRNA165/6 into AGO1 in the endodermis. Given that AGO1 is cell-autonomous, we speculate that *KTN1* suppresses the association of miR165/6 with AGO1 in the endodermis to allow AGO1-unbound miR165/6 to exit the endodermis.

**Figure 5.**
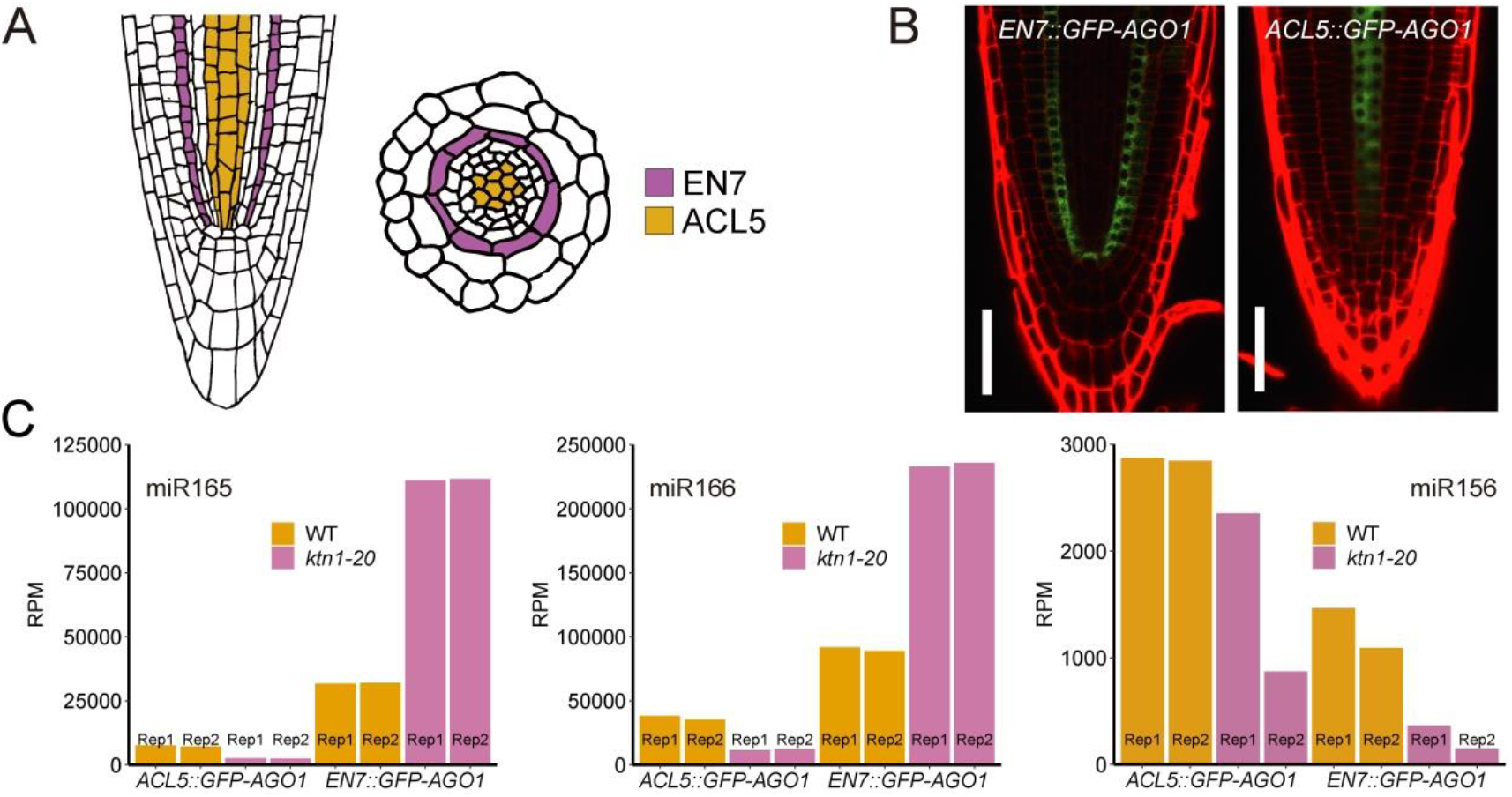
AGO1 loading efficiency of miR165/6 in the endodermis and metaxylem. (A) Schematic representations of a longitudinal (left) and a cross (right) section of an *Arabidopsis* root. Endodermis, and metaxylem and procambia, where *GFP-AGO1* is specifically expressed, are colored. (B) Representative confocal images of EN7::GFP-AGO1 and ACL5::GFP-AGO1. Scale bars, 50 μm. (C) Normalized read counts of miR165, miR166 and miR156 in small RNA-seq of GFP-AGO1 IP from the indicated genotypes. Rep1 and 2 are two independent replicates. See also Figure S6.

### MTs Suppress the Association of miR165/6 with Cytoplasmic AGO1

In the process of miRNA biogenesis, miRNAs are loaded into AGO1 in the nucleus and miRISCs are exported to the cytoplasm (Bologna et al., 2018). We sought to understand how *KTN1* suppresses the loading of miR165/6 into AGO1. We first examined whether *KTN1* affects the nucleo-cytoplasmic partitioning of AGO1, as suppression of AGO1‟s nuclear import could perceivably inhibit miRISC formation during miRNA biogenesis in the nucleus. In *ktn1-20*, the steady-state GFP-AGO1 signals exhibited a diffuse pattern in the cytoplasm similar to those in wild type (Figure S7A). As the steady-state GFP-AGO1 localization did not reveal the nuclear pool of AGO1, we performed nuclear-cytoplasmic fractionation to examine the distribution of AGO1. Results showed that AGO1 protein levels were approximately the same between *ktn1-20* and wild type in either fraction (Figure S7B), indicating that *ktn1-20* did not affect the nuclear-cytoplasmic partitioning of AGO1. Similarly, RNA gel blot assays showed that *ktn1-20* did not alter the levels of eight examined miRNAs or their nucleo-cytoplasmic partitioning (Figure S7C). Thus, *KTN1* does not affect the nuclear import/export of AGO1 or miRISCs.

Given the formation of miRISCs in the nucleus during miRNA biogenesis, for mobile miRNAs, there must be an unloaded fraction that exits the nucleus and then the cell. Upon entry of the next cell, the miRNAs may be loaded into AGO1 or move on to another cell. A major unknown question is whether AGO1 loading occurs in the cytoplasm or the nucleus for mobile miRNAs that arrive in the cytoplasm of a recipient cell. Given that *KTN1* suppresses the loading of miR165/6 into AGO1 without affecting the nucleo-cytoplasmic distribution of AGO1, we entertained the hypothesis that miR165/6 can be loaded into AGO1 in the cytoplasm and *KTN1* suppresses this process. To test this, we fused the ligand-binding domain of mammalian glucocorticoid receptor (GR) to GFP-AGO1 to prevent its nuclear localization (Dittmar et al., 1997; Horstman et al., 2017). *35S::GR-GFP-AGO1* was transiently co-expressed with *35S::MIR165A* and *35S::mScarlet-MAP4* (a marker for MTs) (Pan et al., 2020) in *N. benthamiana* leaves. GR-GFP-AGO1 and mScarlet-MAP4 signals were detected at 36 hours post infiltration. As expected, GR-GFP-AGO1 was exclusively localized in the cytoplasm and mScarlet-MAP4 signals reflected well organized cortical MT arrays (Figure 6A). We next tested whether miR165 could be loaded into cytoplasmic AGO1 and whether MT organization affected cytoplasmic miRISC formation. MTs were disrupted by infiltrating the leaves with the MT depolymerization drug oryzalin at 36 hours after infiltration of the above constructs. After another 24 hours, as indicated by mScarlet-MAP4, MTs were fragmented and depolymerized, while MTs were intact in mock-treated leaves (Figure 6A). We immunoprecipitated GR-GFP-AGO1 and analyzed the levels of miR165 associated with GR-GFP-AGO1 by RNA gel blot analysis. Indeed, miR165 was associated with cytoplasmic GR-GFP-AGO1, indicating that miRISC can form in the cytoplasm. Strikingly, the association of miR165 with GR-GFP-AGO1 was enhanced under oryzalin treatment as compared to mock treatment (Figure 6B). These results indicate that intact and/or dynamic MTs suppress the loading of miR165 into AGO1 in the cytoplasm.

**Figure 6.**
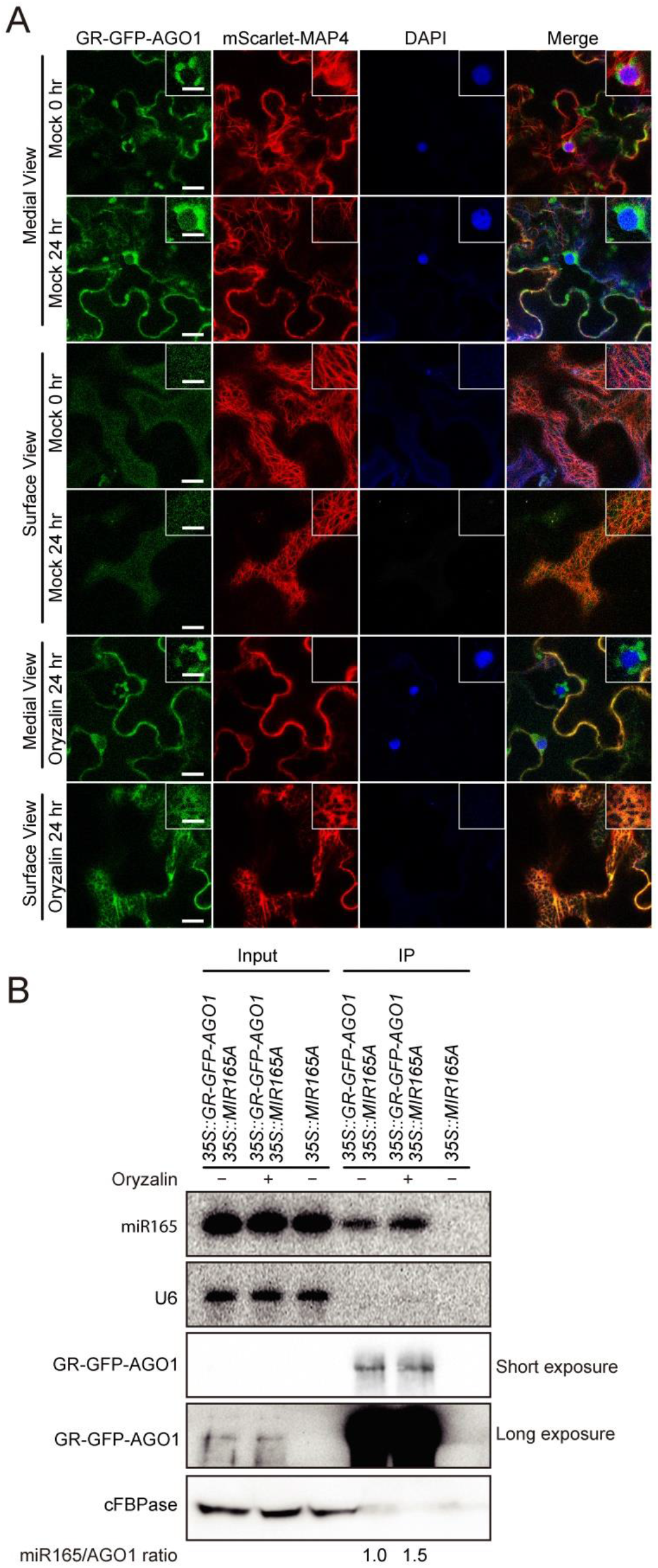
MTs suppress the association of miR165 with AGO1 in the cytosol. (A) Confocal images of GR-GFP-AGO1 and mScarlet-MAP4 co-expressed transiently in *N. benthamiana* leaves. GR-GFP-AGO1 and mScarlet-MAP4 signals were examined before and after oryzalin treatment. Nuclei were stained by DAPI. Note that MTs were disrupted by oryzalin treatment. The medial views show the cytoplasmic localization of GR-GFP-AGO1 and the surface views show patterns of cortical MTs labeled by mScarlet-MAP4. Scale bars, 20 μm. Insets, high magnification images. Scale bar, 10 μm in insets. (B) GR-GFP-AGO1 IP followed by RNA gel blot analysis to examine the association of miR165 with GR-GFP-AGO1 in the cytosol. The levels of miR165 in the immunoprecipitates were quantified against GR-GFP-AGO1 levels and shown below the gel images. *35S::MIR165A* without *GR-GFP-AGO1* was included as a control to show that miR165 signals in the IP samples represented miR165 bound by GR-GFP-AGO1. The longer exposure for the protein gel blot shows the signals of GR-GFP-AGO1 in input samples. See also Figure S7.

## DISCUSSION

In this study, we show that MTs are required for the non-cell autonomous activities of miRNAs with the following lines of evidence: (1) In the *SUC2::amiR-SUL* system, *ktn1* mutations reduce the area of leaf bleaching caused by the mobile amiR-SUL without affecting its accumulation; (2) A *ktn1* mutation abolishes the gradient distribution of PHB controlled by the miR165/6 gradient in the root (Carlsbecker et al., 2010; Miyashima et al., 2011), consequently compromising xylem patterning; (3) A mutation in the MT-associated protein MOR1, similarly affects the non-cell autonomous activities of amiR-SUL and miR165/6 as do *ktn1* mutations. Based on these results, we tentatively conclude that KTN1- and MOR1-regulated microtubule organization is critical to the non-cell autonomous actions of miRNAs.

*KTN1* was previously found to be required for the translation repression activities of miRNAs (Brodersen et al., 2008). Thus, the observed effects of *ktn1* mutations on amiR-SUL or miR165/6 could be explained by *KTN1* aiding their target repression activities rather than their cell-to-cell movement *per se*. However, we were able to show that *KTN1* promotes the cell-to-cell movement of amiR-SUL by measuring its levels in source and recipient tissues (Figure 1J). We also showed that *KTN1* is required in shoots for the shoot-to-root trafficking of amiR-SUL (Figure 1K). Expression of *KTN1* in the endodermis (source of miR165/6) but not the metaxylem (destination of miR165/6) rescued the xylem defects in *ktn1*, while expression of *KTN1* in the protoxylem (tissue miR165/6 transits through) partially rescued the xylem defects. These results collectively suggest that *KTN1* acts in source and intermediary tissues to enable the transport of miRNAs.

In plants, most miRNAs are associated with their effector protein AGO1. By gel filtration, a cytoplasmic pool of AGO1-unbound miRNAs was found (Dalmadi et al., 2019), suggesting that AGO1 loading is not 100% for certain miRNAs. AGO1 is cell-autonomous, as expression of fluorescent protein-tagged AGO1 with layer-specific promoters in the root resulted in corresponding layer-specific signals (Brosnan et al., 2019), which was also found in this study with *EN7::GFP-AGO1* and *ACL5::GFP-AGO1* lines. This indicates that AGO1-unbound miRNAs undergo cell-to-cell movement. In fact, AGO1 was found to “consume” mobile siRNAs as they traverse root cell layers by forming siRISCs to prevent their further movement (Devers et al., 2020). We found that the association of miR165/6 with AGO1 in the endodermis is higher in *ktn1* as compared to wild type, suggesting that *KTN1* suppresses miR165/6-AGO1 RISC formation in the endodermis to enable the exit of AGO1-unbound miR165/6.

In *Arabidopsis*, AGO1 contains a nuclear localization signal and a nuclear export signal that enable its nucleo-cytoplasmic shuttling (Bologna et al., 2018). During miRNA biogenesis, miRNAs are loaded into AGO1 in the nucleus and exported into the cytoplasm as miRISCs (Bologna et al., 2018). Some siRNAs, such tasiRNAs, are produced and loaded into AGO1 in the cytosol (Bologna et al., 2018). We rationalize that mobile miRNAs are the fractions that escape AGO1 loading in the nucleus and enter the cytosol. Given the presence of cytoplasmic AGO1, the miRNAs must avoid being loaded into AGO1 in the cytoplasm to exit the cell. Our studies with cytoplasmically sequestered AGO1 indicate that miR165/6 can be loaded into AGO1 in the cytoplasm and disruption of MTs enhances this loading. Thus, we propose that cytoplasmic miRISC formation needs to be suppressed in source cells to enable miRNA‟s cell-to-cell movement and MTs play a role in this process (Figure 7). In fact, during transit, cytoplasmic RISC formation would limit the range of miRNA‟s cell-to-cell movement, which is consistent with the observation that expression of *KTN1* in the protoxylem partially rescues the *ktn1* mutant phenotypes. How MTs suppress the cytoplasmic loading of miRNAs is unknown. Intriguingly, we found co-localization of AGO1 and MTs, particularly when oryzalin was applied (Figure 6A). Perhaps a fraction of AGO1 proteins normally associates with/dissociates from MTs in a dynamic manner, and the dissociation is attenuated when MTs are disrupted. If this is the case, a direct role of MTs in regulating the cytoplasmic loading of miRNAs is possible.

**Figure 7.**
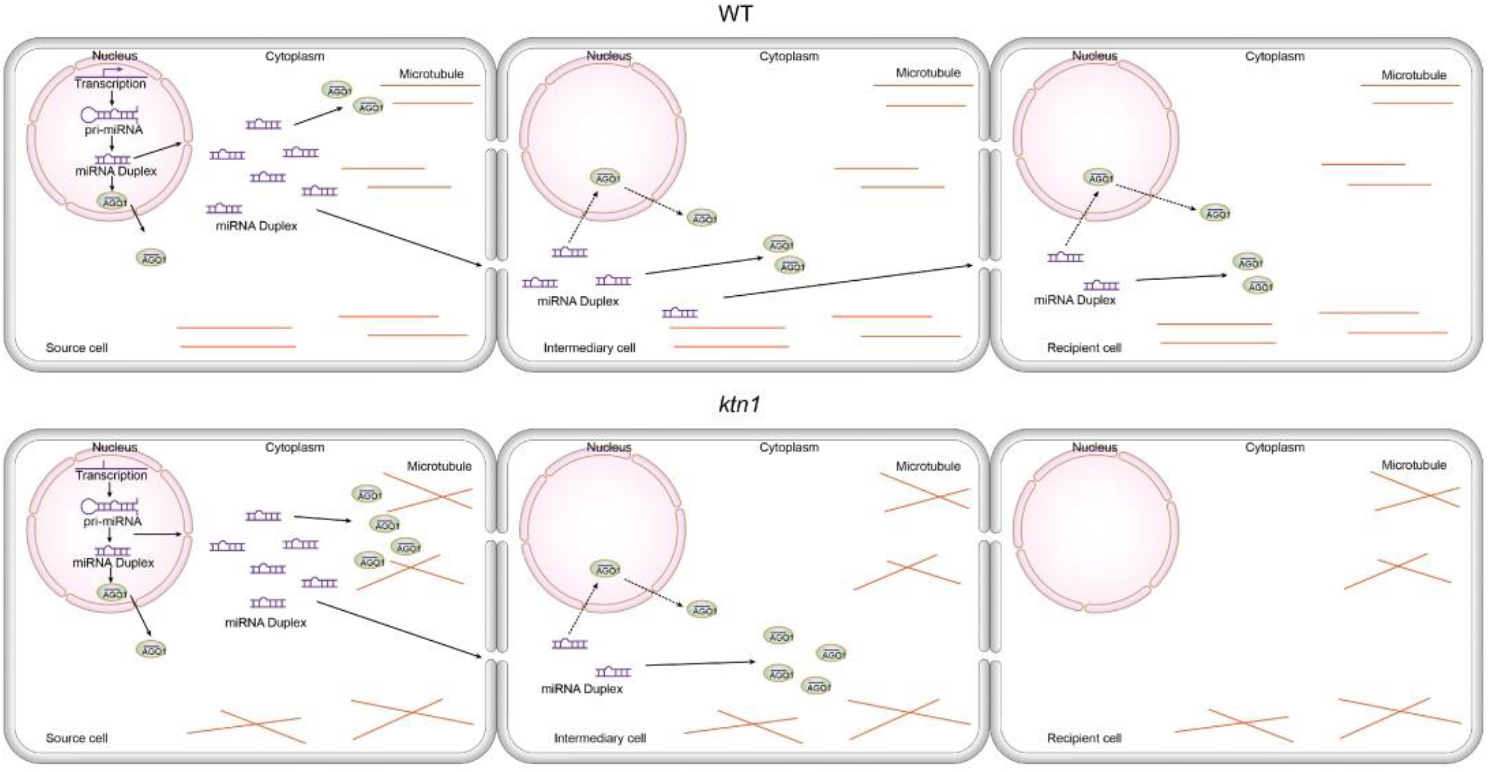
Model of MTs promoting the cell-to-cell movement of miRNAs by suppressing their cytoplasmic AGO1 loading. MiRNA biogenesis occurs in the nucleus in the source cell (left) in both WT and *ktn1* mutants. A portion of the miRNA/miRNA* duplexes exits the nucleus. In WT, organized and/or dynamic MTs suppress the association of miRNAs with the cell-autonomous AGO1 in the cytosol in the source and intermediary cells, allowing miRNAs to exit the cells. In *ktn1* mutants, miRNA/miRNA* duplexes are loaded into AGO1 in the cytoplasm in the source cell, resulting in a lower level of duplexes exiting the cell. In the intermediary cell, the duplexes are loaded into AGO1 in the cytoplasm, leaving none to arrive at the destination. Orange lines represent MTs. Dotted arrows indicate that it is unknown whether mobile miRNA/miRNA* duplexes also enter the nucleus to be loaded into AGO1 in intermediary and recipient cells. It should also be noted that it is unknown whether miRNA/miRNA* duplexes are the mobile agents that exit the nucleus or the source cell; single-stranded, AGO1-unbound miRNAs can also be potential mobile agents.

*KTN1* was shown to be required for miRNA-mediated translational repression in plants (Brodersen et al., 2008). In this study, we uncovered a role of *KTN1* in promoting the intercellular movement of miRNAs. Intriguingly, in the *ktn1-20* mutant, the target of amiR-SUL was de-repressed at the protein but not mRNA level, consistent with the requirement of *KTN1* for amiR-SUL‟s translation repression activity. Are the functions of *KTN1* in mediating translation repression by miRNAs and promoting the intercellular movement of miRNAs related or independent? The answer awaits future research.

## Supporting information

Supplemental Information includes seven figures and one table.

## ACKNOWLEDGMENTS

We thank Chenjiang You and Brandon H. Le for training in data analysis. We are grateful to Drs. Detlef Weigel, Zhenbiao Yang and Venugopala (Reddy) Gonehal for sharing the *SUC2::amiR-SUL* transgenic line, the *UBQ10::mScarlet_MAP4* plasmid and the *pGreen-WUS-GR* plasmid, respectively. We thank Brandon H. Le for comments on the manuscript. This work was supported by National Institutes of Health (GM129373) to X.C.

## AUTHOR CONTRIBUTIONS

X.C. and L.F. designed the experiments. X.C. supervised the project. L.F., C.Z. and Y.Z. performed the experiments. L.F., E.S. and J.J. analyzed the data. X.C., L.F. and K.N. wrote the manuscript.

## METHODS

### Plant materials and growth conditions

*Arabidopsis thaliana* strains used in this study were all in the Columbia-0 (Col-0) background. Seeds were sterilized and grown on half-strength MS medium (PH 5.7) under long-day conditions (16 h light/8 h dark). The *SUC2::amiR-SUL* line was from Detlef Weigel (Max Planck Institute for Developmental Biology, Tubingen, Germany) (de Felippes et al., 2011). *ktn1-20* was a new allele isolated from an EMS mutagenesis screen in the *SUC2::amiR-SUL* background. *ktn1-2* (SAIL343_D12) is a T-DNA insertion mutant. Transgenic lines of *PHB::PHB-GFP, pMIR165A::GFPer, pMIR165B::GFPer, pMIR166A::GFPer, pMIR166B::GFPer* and the plasmid of *CRE1::MIR165A* were described (Miyashima et al., 2011). *UBQ10::mScarlet_MAP4* was a gift from Drs. Jingzhe Guo and Zhenbiao Yang (Pan et al., 2020).

### Mutagenesis and Screening

Seeds of a homozygous *SUC2::amiR-SUL* transgenic line were subjected to ethyl methanesulfonate mutagenesis as described (Jia et al., 2017; Zhang et al., 2020). The *ktn1-20* mutant was isolated based on its weaker leaf bleaching phenotype. The genomic sequence containing the coding region of *KTN1* was amplified by PCR with primers KTN1-seq-F and KTN1-seq-R and the PCR product was subjected to sequencing to identify the mutation. See Table S1 for sequences of oligonucleotides.

### Plasmid Construction and Plant Transformation

All constructs were made in a modified pCambia1300 vector, which contains the 35S promoter inserted between the *Eco*RI and *Sac*I sites and the NOS terminator inserted between the *Pst*I and *Hind*III sites. For genetic complementation, a genomic fragment of *KTN1*, which contains the promoter and coding regions, were amplified from *Arabidopsis* genomic DNA using primers KTN1g-F and KTN1g-R and recombined into the modified pCambia1300 that was digested with *Eco*RI and *Sal*I to remove the 35S promoter. To create plasmids for root cell-layer-specific expression of *GFP-AGO1*, *EN7* and *ACL5* promoters were amplified using primers EN7-F/EN7-R and ACL5-F/ACL5-R, respectively, from *Arabidopsis* genomic DNA and inserted into pCambia1300 to replace the 35S promoter, resulting in pCambia1300-EN7 and pCambia1300-ACL5, respectively, and then *GFP* and *AGO1* genomic fragments were PCR-amplified with primers GFP-F/GFP-R and AGO1-F/AGO1-R, respectively, and recombined downstream of the *EN7* and *ACL5* promoters. Constructs for root layer-specific *KTN1* expression were obtained by inserting *EN7*, *ACL5*, *AHP6* and *SHR* promoters into pCambia1300 and then cloning the genomic fragment containing the coding region of *KTN1* downstream of the root layer-specific promoter. The primers for *EN7*, *ACL5*, *AHP6*, *SHR*, and *KTN1* PCR were EN7-F/EN7-R, ACL5-F/ACL5-R, AHP6-F/AHP6-R, SHR-F/SHR-R, and KTN1-coding-F/KTN1g-R, respectively. To construct *35S::GR-GFP-AGO1*, the ligand binding domain of *GR* was PCR-amplified from the *pGreen-WUS-GR* plasmid (a gift from Venugopala (Reddy) Gonehal) (Yadav et al., 2010) using primers GR-F and GR-R and inserted into pCambia1300. *GFP* and an *AGO1* genomic fragment were PCR-amplified with primers GFP-F/GFP-R(GR) and AGO1-F(GR)/AGO1-R(GR), respectively, and fused with GR. *AGO1::GFP-AGO1* was constructed as follows. A genomic fragment containing the *AGO1* promoter was PCR-amplified using the primer pair AGO1p-F/AGO1p-R and inserted into pCambia1300 and then *GFP* and *AGO1* were PCR-amplified using primers pairs GFP-F/GFP-R and AGO1-F/AGO1-R, respectively and inserted downstream of the *AGO1* promoter. *35S::MIR165A* was constructed by inserting a 300 bp genomic fragment containing the pri-miR165A region (amplified by PCR with primers MIR165A-F and MIR-165A-R) downstream of the 35S promoter in pCambia1300. All constructs were introduced into *Agrobacterium tumefaciens* (*A. tumefaciens*) by electroporation. Stable transgenic plants were produced through *Agrobacterium tumefaciens*-mediated floral dip transformation (Clough and Bent, 1998).

### Quantification of the Leaf Bleaching Phenotype

Color images of individual leaves were obtained with an Epson scanner. Image analyses were carried out in Python 3.6.5 using the open-source Open CV package (Bradski and Kaehler, 2000). Leaves were segmented from the background using a color threshold in the L*a*b* color space. From the thresholded images, binary masks were produced for each leaf. A mean blur with a kernal size of 3×3 was applied to each leaf to remove noise. The area of bleached veins was determined using an adaptive Gaussian threshold with a block size of 101 pixels. Binary masks of the bleached vein area were produced for each leaf. Binary masks for the inter-vein regions were calculated by subtracting the vein masks from the leaf masks. Leaf area, vein area and inter-vein area were calculated by summing the pixel values of the respective binary masks. To quantify the extent of bleaching, color images were transformed from RGB to HSV (hue, saturation, value) and the hue channel was extracted. Hue values range from 0 to 360 with yellow having a value of 60 and green having a value of 120. The mean values for the entire leaf, bleached vein area and inter-vein area were calculated with the NumPy package (Oliphant, 2006) using the respective binary masks for the original image. All values were written to a single csv file for further analysis.

### Epidermal, Mesophyll and Vascular Tissue Separation

The separation of epidermal, mesophyll and vascular tissues was performed according to the Meselect method (Svozil et al., 2016). Briefly, three-week-old rosette leaves were placed between two tape strips. After gently peeling off the two tapes, the epidermal tissues on the abaxial side of the leaves were separated from the vascular tissue and the adaxial side of the leaves. The tapes with the abaxial epidermis were incubated in protoplasting solution (1% cellulase Onozuka R10 (Yakult), 0.25% Maceroenzyme R10 (Yakult), 0.4 M mannitol, 10 mM CaCl_2_, 20 mM KCl, 0.1% bovine serum albumin, 20 mM MES (PH5.7)) for 15 min at room temperature on a shaker at 50 rpm to remove the residual mesophyll cells and then washed twice with washing buffer (154 mM NaCl, 125 mM CaCl_2_, 5 mM KCl, 2 mM MES, pH 5.7). The tapes with the abaxial epidermis were collected as epidermal tissue. The other tapes with the remaining leaf tissues were incubated in protoplasting solution for 45 min. The enzyme solution containing mesophyll cells was collected and centrifuged at 100g for 5 min at 4°C. The pellet was washed twice with washing buffer and collected as mesophyll cells. The vascular tissue was removed from the tapes using forceps and washed twice with washing buffer. The epidermal tapes, mesophyll cells and vascular tissue were frozen in liquid nitrogen and store at - 80°C before RNA extraction.

### Quantitative RT-PCR

Total RNA was treated with DNase I (Sigma) at 37°C for 1 hour and reverse transcription was performed with RevertAid Reverse Transcriptase (ThermoFisher Scientific) using an oligo(dT) primer. Quantitative PCR was performed on the BioRad CFX96 system with SYBRGreen Supermix (BioRad). Relative expression levels were calculated using the pcr package in R (Ahmed and Kim, 2018). The *UBQ5* transcript was detected in parallel and used for normalization. The primers used are listed in Table S1.

### Grafting

Grafting was performed according to the method described (Marsch-Martínez et al., 2013). Briefly, plants were grown vertically on ½ MS medium under short-day conditions (8 h light/16 h dark). 7-day-old seedlings were used for grafting. Seedlings were placed on a thin layer of 1% agarose and cotyledons were removed from the scion. Hypocotyls were cut in the middle and scions and root stocks were aligned under a binocular stereoscope. The grafted plants were transferred back to the short-day growth chamber and incubated vertically for another 10 days. Shoots and roots were collected separately from 20-30 seedlings and subjected to RNA extraction using TRI reagent.

### Microscopy and Imaging

For xylem phenotype analysis, 5-day-old seedlings were stained in 0.0001% basic fuchsin in 95% ethanol for 5 min, and then washed two times with 70% ethanol (Smith et al., 2013). Primary roots were cut and mounted in 50% glycerol, and observed under an Olympus BX53 fluorescent microscope. To quantify the distribution pattern of PHB-GFP, 10-15 individual roots were imaged under confocal laser-scanning microscopy (CLSM). Fluorescent signals across the root were quantified using ImageJ. CLSM was performed on Leica SP5 and Zeiss LSM 780 microscopes. Roots were stained with 10 μM propidium iodide followed by CLSM.

### Nuclear-cytoplasmic Fractionation

12-day-old seedlings were ground into fine powder in liquid nitrogen and the powder was resuspended with lysis buffer (20 mM Tris-HCl at PH 7.5, 20 mM KCl, 2 mM EDTA, 2.5 mM MgCl_2_, 25% glycerol, 250 mM Suc, 5 mM DTT and proteinase inhibitor cocktail (Roche)) at 2ml/g. The suspension was filtered through double layers of Miracloth and centrifuged at 1500g for 15 min. The supernatant was transferred into a new tube and centrifuged at 10,000g for 10 min at 4°C. The supernatant was collected as the cytoplasmic fraction. The pellet was then washed six times with nuclear resuspension buffer (NRB: 20 mM Tris-HCl, pH 7.4, 25% glycerol, 2.5 mM MgCl_2_, and 0.2% Triton X-100). The pellet was then resuspended with 500 μl NRB2 (20 mM Tris-HCl, pH 7.5, 0.25 M sucrose, 10 mM MgCl_2_, 0.5% Triton X-100, and 5 mM 2-mercaptoethanol) and the suspension was carefully laid on top of 500 μl NRB3 (20 mM Tris-HCl, pH7.5, 10 mM MgCl_2_, 1.7 M sucrose, 0.5% Triton X-100, 5 mM 2-mercaptoethanol and protease inhibitor cocktail (Roche). The samples were centrifuged at 16,000g for 45 min at 4°C. The pellet was collected as the nuclear fraction. 10% of the cytoplasmic and nuclear fractions were boiled in 1 X SDS sample loading buffer (50 mM Tris-Cl at pH 6.8, 2% SDS, 0.1% bromophenol blue, 10% glycerol, 1% 2-mercaptoethanol) and saved for protein gel blot analysis. The remaining cytoplasmic and nuclear fractions were subjected to RNA extraction using TRI reagent.

### RNA Gel Blot Analysis of Small RNAs

12-day-old seedlings were ground into powder in liquid nitrogen and RNA was extracted using the TRI reagent (Molecular Research Center, Inc) according to the manufacturer‟s instructions. Small RNA gel blot analysis was performed as described (Li et al., 2016). 5 μg total RNA from Col and *ktn1-20* was resolved on 15% Urea-PAGE gels and transferred onto Hybond NX membranes (GE Healthcare Life Sciences) followed by chemical cross-linking using EDC (1-ethyl-3-(3-dimethylaminopropyl) carbodiimide hydrochloride). Complementary oligonucleotides were end-labeled with ^32^P and used to probe the membrane. The bands on the RNA gel blot were quantified by ImageQuant TL 8.1 and normalized against U6 or tRNA. The oligonucleotide probes used are listed in Table S1.

### Immunoprecipitation of Root Layer-specific GFP-AGO1

9-day-old *Arabidopsis* roots of *EN7::GFP-AGO1* and *ACL5::GFP-AGO1* transgenic lines in WT and *ktn1-20* backgrounds were collected and ground into powder in liquid nitrogen. Cells were lysed in immunoprecipitation buffer (50 mM Tris 7.5, 150 mM NaCl, 4mM MgCl_2_, 2mM DTT, 10% Glycerol, 0.1% NP-40, 1x proteinase inhibitor cocktail (Roche)) by gentle rotation at 4°C for 30 min. Cell debris was removed by centrifugation twice at 15000 rpm for 20 min at 4°C. The supernatant was incubated with GFP-Trap_MA (ChromoTek) for 2 h with gentle rotation. The beads were washed 5 times with washing buffer (50 mM Tris 7.5, 150 mM NaCl, 4mM MgCl_2_, 2mM DTT, 10% Glycerol, 0.5% NP-40, 1x proteinase inhibitor cocktail (Roche). Beads were magnetically collected and 10 % GFP-AGO1 immunoprecipitates were boiled in 1X SDS sample loading buffer for protein gel blot analysis and the remainder was subjected to RNA extraction using the TRI reagent.

### Protein Gel Blot Analysis

Total proteins from twelve-day-old seedlings were resolved by SDS-PAGE, transferred to nitrocellulose membranes, blocked in 1X PBS + 0.05% Tween-20 supplemented with 5% non-fat milk for 1 hr and incubated with primary antibodies at 4℃ overnight. After washing three times in 1X PBS + 0.05% Tween-20, membranes were incubated with horseradish peroxidase-conjugated secondary antibodies for 1 hour, then rinsed three times and detected with ECL western blotting detection reagent (Cytiva Amersham).

### Small RNA Sequencing and Data Analysis

To construct small RNA libraries from total RNA, 20 μg total RNA from 14-day-old seedlings was resolved in a 15% Urea-PAGE gel and small RNAs of 15-40 nt were isolated from the gel and subjected to small RNA library construction using NEBNext Multiplex Small RNA Library Prep Set for Illumina (E7300). Small RNAs extracted from GFP-AGO1 IP products were processed into sequencing libraries using NEBNext Multiplex Small RNA Library Prep Set for Illumina (E7300) without fractionation by gel electrophoresis. The libraries were sequenced on an Illumina HiSeq 2500. sRNA-seq data were analyzed using the pRNASeqTools pipeline (https://github.com/grubbybio/pRNASeqTools). Raw reads from sRNA-seq were first trimmed to remove the adaptor sequence (AGATCGGAAGAGC) by cutadapt 3.0. The trimmed reads were mapped to the *Arabidopsis* genome (Araport 11) using Shortstack 3.4 with parameter “‘--bowtie_m 1000 --ranmax 50 --mmap u --mismatches 0”. The miRNA levels were quantified by calculating the RPM (reads per million mapped reads). Two biological replicates for each genotype were included.

### Transient Expression of GR-GFP-AGO1 and IP in N. benthamiana

Transient expression in *N. benthamiana* was performed as described (Martin et al., 2009). The *A. tumefaciens* strain GV3101 carrying *35S::MIR165A*, *35S::GR-GFP-AGO1* and *UBQ10::mScarlet-MAP4* were infiltrated into the leaves of three-week-old *N. benthamiana*. At 36 hours post infiltration, the leaves were infiltrated with 20 μM oryzalin (Sigma) or 0.02% DMSO (mock treatment). Leaves were collected at 24 hours post oryzalin or mock infiltration. IP was performed using GFP-Trap_MA (ChromoTek) as described above. The imaging of GR-GFP-AGO1 and mScarlet-MAP4 was performed using CLSM as described above. 5 ug/ml DAPI was used to stain the nuclei.

## Data and Material Availability

All sequencing datasets reported in this paper have been deposited in the NCBI database under the accession number GSE168805.

