## Supplemental Information includes seven figures and one table. for "Microtubules Promote the Non-cell Autonomy of MicroRNAs by Inhibiting their Cytoplasmic Loading into ARGONAUTE1 in *Arabidopsis*"

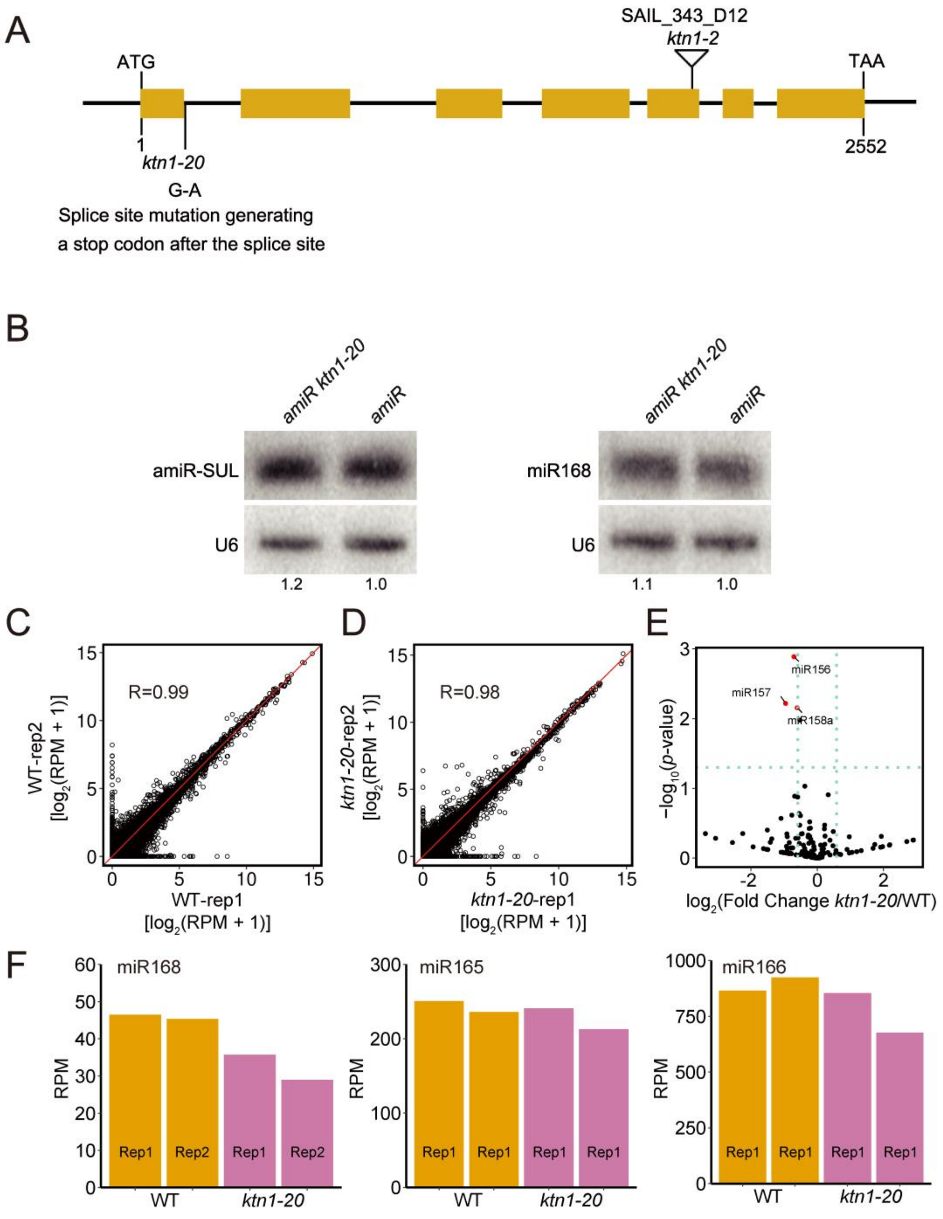


**Figure S1.** ***KTN1*** **Is not Required for the Biogenesis of miRNAs, related to Figure 1**

(A) A diagram of the *KTN1* gene structure showing the mutation sites of *ktn1-20* and *ktn1-2*. Rectangles represent exons. (B) RNA gel blot assays showing the relative levels of amiR-SUL and miR168 in *SUC2::amiR-SUL (amiR)* and *amiR ktn1-20.* U6 was the loading control. (C, D) Scatter plots showing the correlation between two biological replicates of small RNA-seq for each genotype (WT and *ktn1-20*). Normalized read counts in 100-bp bins were used for Pearson correlation analysis. (E) A volcano plot showing differentially expressed miRNAs in *ktn1-20* vs. WT (fold change > 1.5, *p*-value < 0.05). Only miR156, miR157 and miR158a show significant reductions in *ktn1-20* vs. WT. miR156 represents all reads from any of the six *MIR156A-F* genes. miR157 represents all reads from any of the three *MIR157A-C* genes. (F) Normalized read counts of miR165, miR166 and miR168 in small RNA-seq in WT and *ktn1-20*. The two bars for each genotype represent two biological replicates.

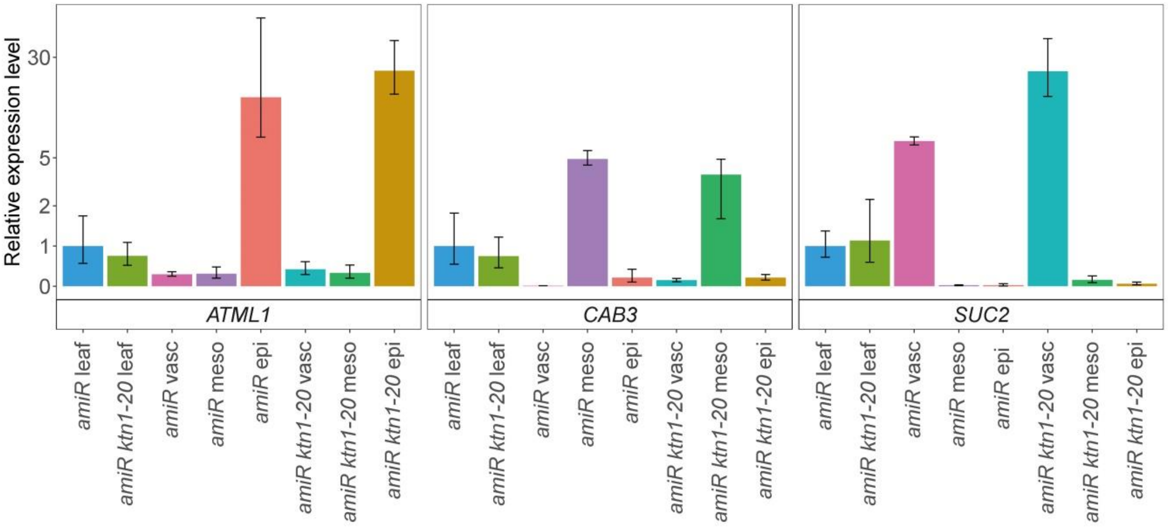


**Figure S2. Quantitative RT-PCR analysis of *ATML1*, *CAB3* and *SUC2* Transcripts, related to Figure 1**

Levels of these transcripts were determined in either whole leaves or Meselect-separated vasculature (vasc), mesophyll tissue (meso) and abaxial epidermal tissue (epi) in *SUC2::amiR-SUL* (*amiR*) and *amiR ktn1-20*. Transcript levels were normalized to those of *UBQ5*. Error bars represent lower and upper interval of the normalized relative transcript levels from three technical replicates.

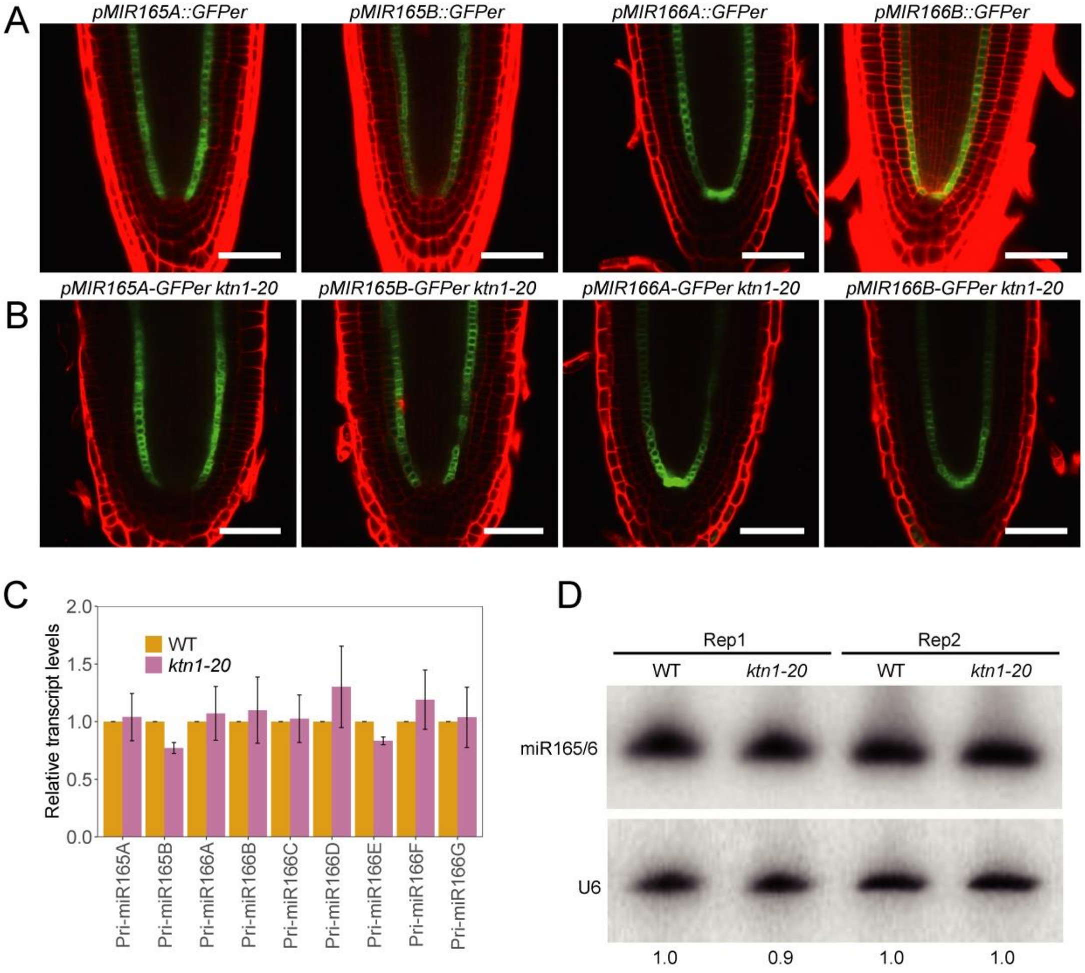


**Figure S3. *KTN1* Is not Required for the Biogenesis of miR165/6, related to Figure 2.**

(A, B) Promoter activities of *MIR165A*, *MIR165B*, *MIR166A* and *MIR166B* in WT and *ktn1-20* as shown by the expression of the GFPer reporter driven by the respective promoters. Scale bars, 50 µm. (C) Levels of various pri-miR165 and pri-miR166 transcripts in WT and *ktn1-20* as determined by quantitative RT-PCR. *UBQ5* RNA served as the internal control. Error bars represent SD calculated from three biological replicates. (D) RNA gel blot analysis of miR165/6 in WT and *ktn1-20*. U6 was used as a loading control. Two biological replicates (Rep) are shown. The numbers below the gel images show relative levels.

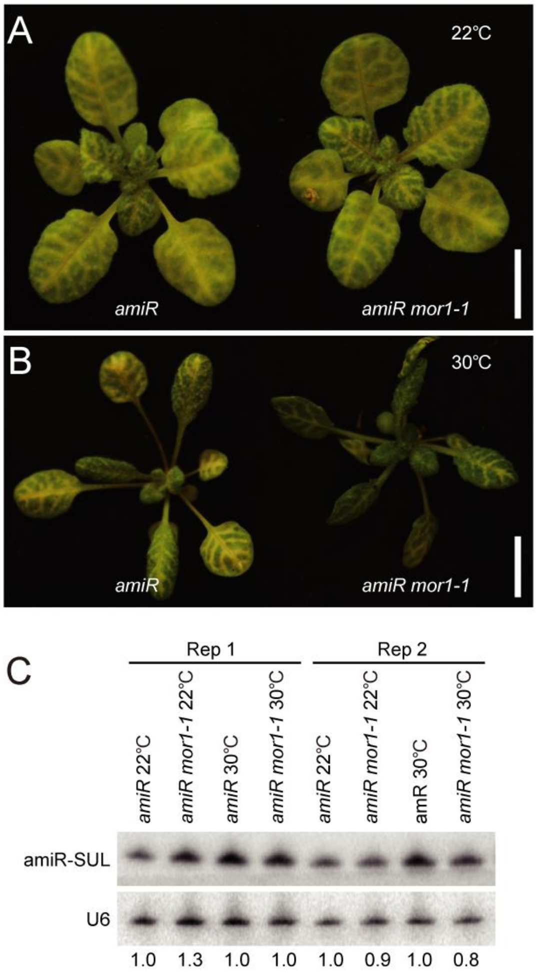


**Figure S4. Effects of the *mor1-1* Mutation on *SUC2::amiR-SUL* Plants, related to Figure 2.**

(A) Three-week-old plants of *SUC2::amiR-SUL* (*amiR*) and *amiR mor1-1* grown at 22℃. No obvious differences in vein-centered bleaching between the two genotypes were observed. Scale bars, 1cm. (B) 10-day-old plants of *amiR* and *amiR mor1-1* grown at 22℃ were transferred to 30℃ and grown for another 10 days. *amiR mor1-1* plants exhibited weaker leaf chlorosis. Scale bars, 1cm. (C) RNA gel blot analysis of amiR-SUL in *amiR* and *amiR mor1-1* grown under either 22℃ or 30℃ as described above. Relative signal intensities were normalized against U6 and values were given. Two biological replicates (Rep) were included.

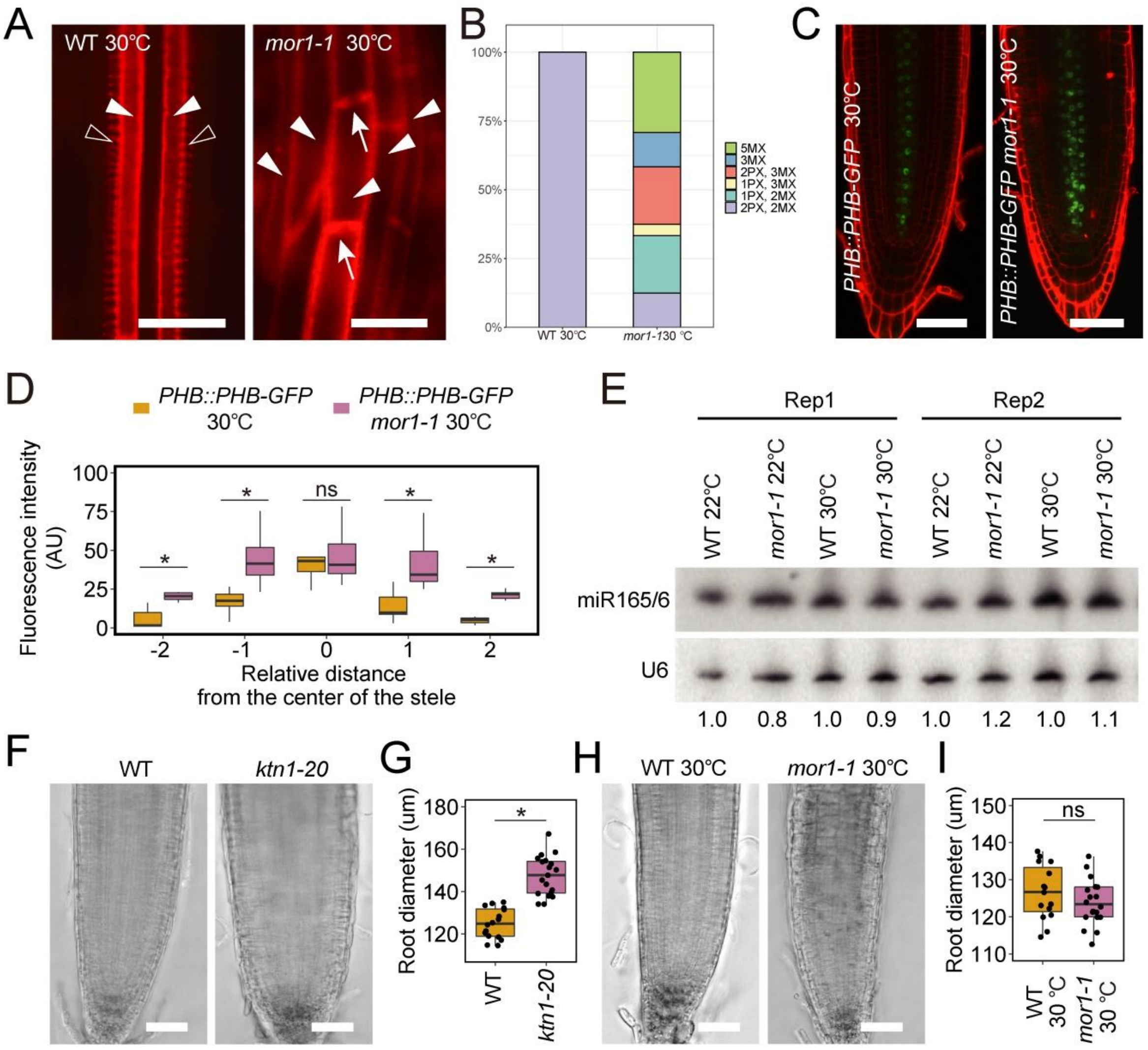


**Figure S5. Xylem Patterns and PHB-GFP Distribution in *mor1-1*, related to Figure 2.**

(A) Representative images showing xylem patterns in WT and *mor1-1* roots at 30℃. Protoxylem is less recognizable in *mor1-1* roots. Unfilled arrowheads indicate protoxylem; filled arrowheads indicate metaxylem. Note the cell wall (arrows) between two adjacent cells, a feature of metaxylem, in *mor1-1*. Scale bars, 20 µm. (B) Quantification of xylem phenotypes in WT (n = 23) and *mor1-1* (n = 24) at 30℃. (C) PHB-GFP distribution in WT and *mor1-1* at 30℃. Scale bars,

50 µm. (D) Quantification of PHB-GFP signal intensity in WT (n = 10) and *mor1-1* (n = 14) roots at 30℃. AU: arbitrary unit. (E) RNA gel blot analysis of miR165/6 in WT and *mor1-1* at either 22℃ or 30℃. U6 was used as a loading control. The numbers indicate relative levels of amiR-SUL with WT set as 1.0. Two biological replicates (Rep) are shown. (F) Representative images of WT and *ktn1-20* roots. (G) Quantification of root diameter in WT and *ktn1-20* at 400 µm from the root tip. (H) Representative images of WT and *mor1-1* roots at 16 hours after plants were transferred to 30℃. (I) Quantification of root diameter in WT and *mor1-1* at 16 hours after plants were transferred to 30℃. Measurements were taken at 400 µm from the root tip. 15-20 roots were included for quantification. *, *p*-value < 0.01; ns, not significant.

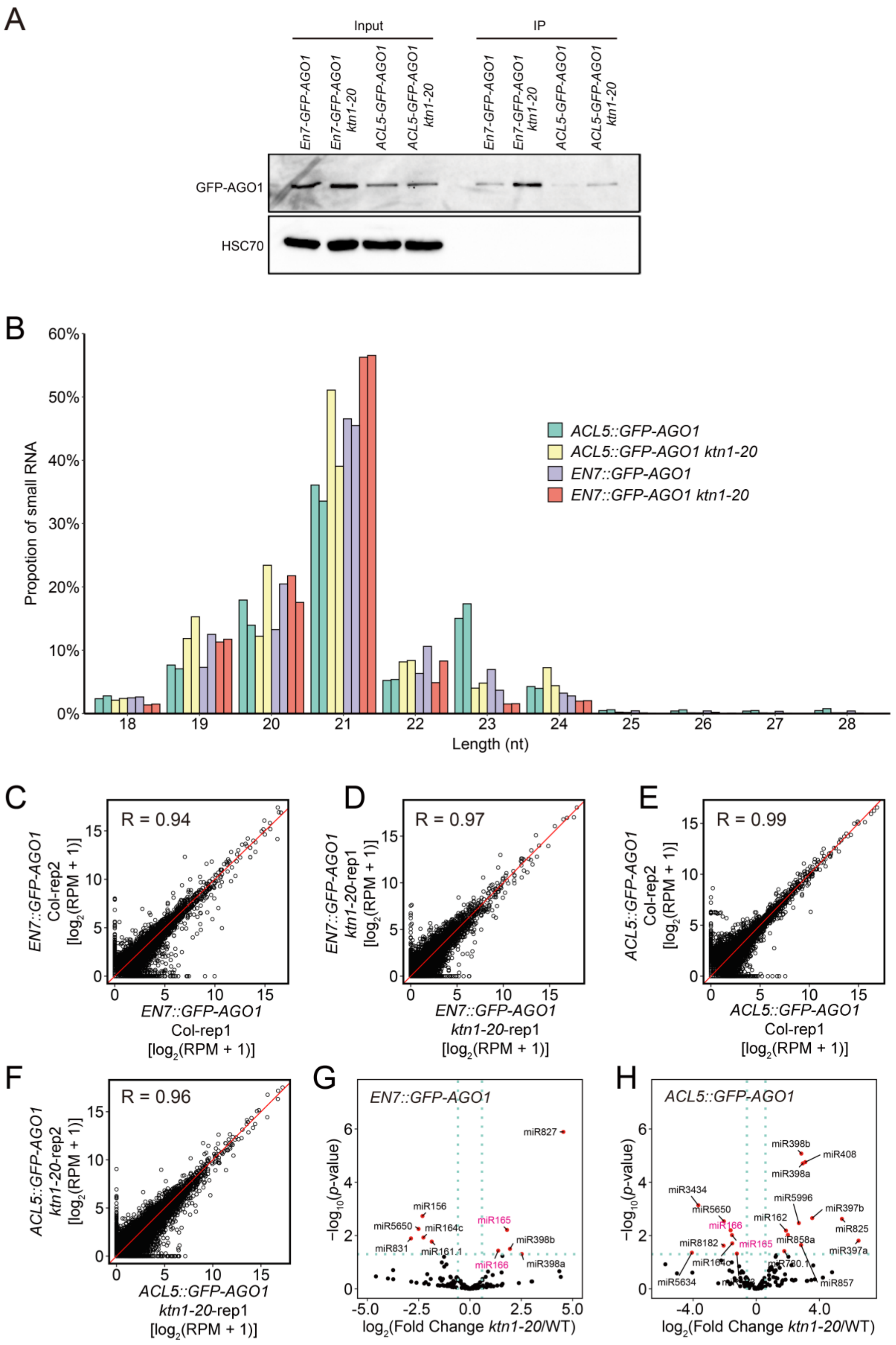
A
Input
IP
En7-GFP-AGO1
En7-GFP-AGO1 ktn1-20
ACL5-GFP-AGO1
ACL5-GFP-AGO1 ktn1-20
GFP-AGO1
HSC70
B
ACL5::GFP-AGO1
ACL5::GFP-AGO1 ktn1-20
EN7::GFP-AGO1
EN7::GFP-AGO1 ktn1-20
Propotion of small RNA
Length (nt)
C
R = 0.94
EN7::GFP-AGO1 Col-rep2 [log2(RPM + 1)]
EN7::GFP-AGO1 Col-rep1 [log2(RPM + 1)]
D
R = 0.97
EN7::GFP-AGO1 ktn1-20-rep1 [log2(RPM + 1)]
EN7::GFP-AGO1 ktn1-20-rep1 [log2(RPM + 1)]
E
R = 0.99
ACL5::GFP-AGO1 Col-rep2 [log2(RPM + 1)]
ACL5::GFP-AGO1 Col-rep1 [log2(RPM + 1)]
F
R = 0.96
ACL5::GFP-AGO1 ktn1-20-rep2 [log2(RPM + 1)]
ACL5::GFP-AGO1 ktn1-20-rep1 [log2(RPM + 1)]
G
EN7::GFP-AGO1
miR827
miR156
miR165
miR5650
miR164c
miR398b
miR831
miR161.1
miR166
miR398a
−log10(p-value)
log2(Fold Change ktn1-20/WT)
H
ACL5::GFP-AGO1
miR398b
miR408
miR398a
miR3434
miR5996
miR5650
miR397b
miR166
miR162
miR825
miR8182
miR165
miR858a
miR397a
miR164c
miR5634
miR857
−log10(p-value)
log2(Fold Change ktn1-20/WT)

**Figure S6. GFP-AGO1 Immunoprecipitation from Endodermis and Metaxylem Followed by Small RNA-seq, related to Figure 5.**

(A) Protein gel blot analysis of GFP-AGO1 from input and immunoprecipitates (IP) from the indicated genotypes. (B) Length distribution of mapped reads from small RNA-seq of GFP-AGO1 IP products from the indicated genotypes. The two bars of the same color represent two biological replicates. (C-F) Scatter plots showing the correlation between two biological replicates of small RNA-seq performed on GFP-AGO1 immunoprecipitates from *EN7::GFP-AGO1* (C), *EN7::GFP-AGO1 ktn1-20* (D), *ACL5::GFP-AGO1* (E) and *ACL5::GFP-AGO1 ktn1-20* (F). Normalized read counts in 100-bp bins were used for Pearson correlation analysis. Pearson correlation coefficients of the two biological replicates are shown in each plot. (G-H) Volcano plots showing differentially AGO1-loaded miRNAs between WT and *ktn1-20* in *EN7::GFP-AGO1* (G) and *ACL5::GFP-AGO1* (H) with fold change > 1.5 and *p*-value < 0.05. All differentially AGO1-loaded miRNAs are represented by red dots. miR165 and miR166 are colored in magenta to show the enhanced AGO1 loading in the endodermis and decreased AGO1 loading in the metaxylem in *ktn1-20* relative to WT.

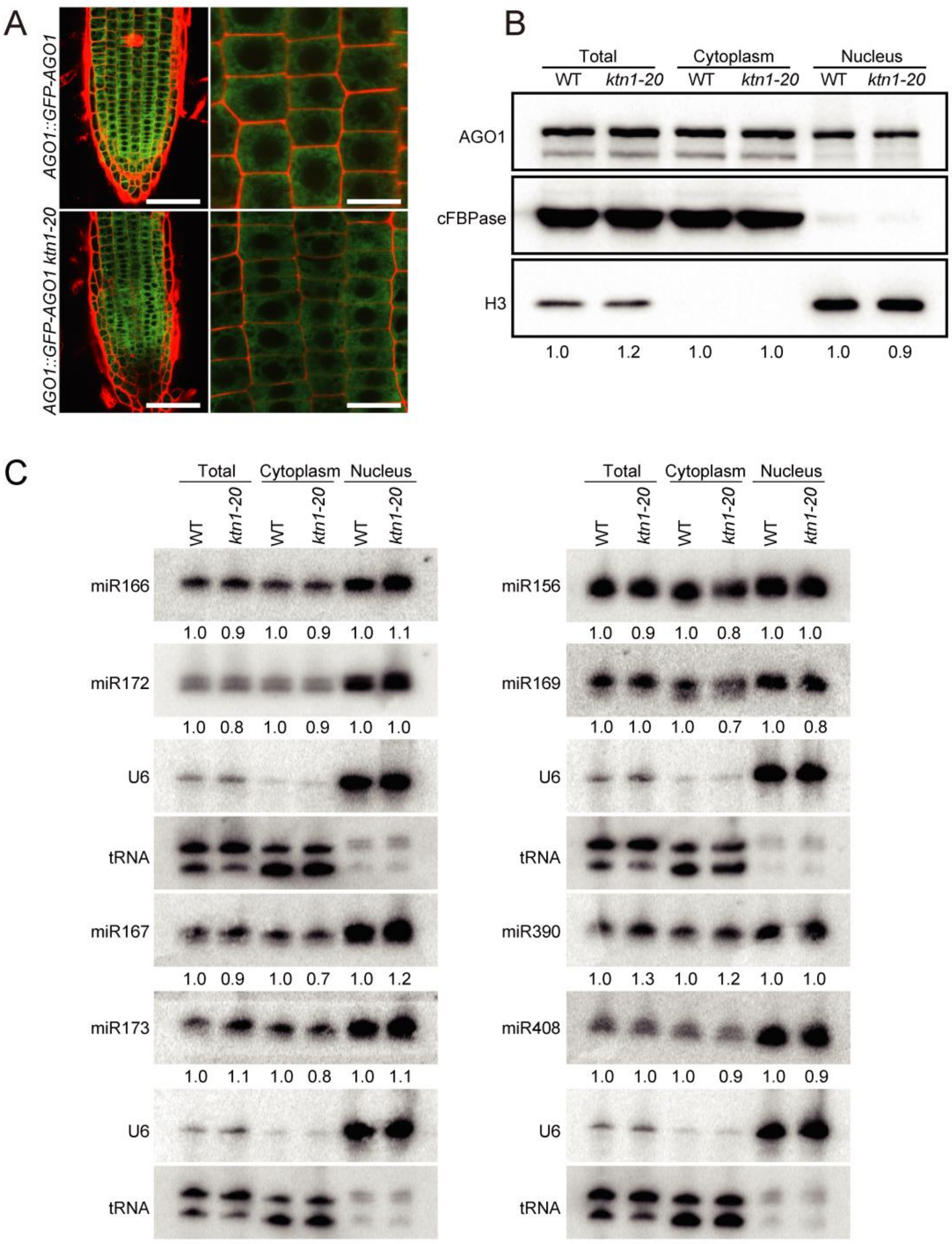


**Figure S7. Subcellular Localization of AGO1 and miRNAs, related to Figure 6.**

(A) Confocal images of GFP-AGO1 in WT and *ktn1-20.* Left and right panels represent low and high magnification, respectively. Green, GFP-AGO; red, propidium iodide (PI) staining showing the outlines of the cells. Scale bars, 50 µm (left); 20 µm (right). (B) Protein gel blot analysis showing the nucleo-cytoplasmic partitioning of AGO1. H3 serves as a nuclear marker, and cFBPase serves as a cytoplasmic marker. AGO1 levels in the cytoplasm and the nucleus were normalized to cFBPase and H3, respectively. AGO1 levels in *ktn1-20* relative to WT are indicated by the numbers below the gel images. (C) RNA gel blot analyses of miRNAs from total extracts (Total), the cytoplasmic fraction (Cytoplasm) and the nuclear fraction (Nucleus) in WT and *ktn1-20*. U6 and tRNA were used as loading controls for nuclear and cytoplasmic RNAs, respectively. Levels in *ktn1-20* relative to WT are indicated by the numbers below the gel images

.

**Supplementary Table 1. DNA Oligonucleotides, Related to Experimental Procedures.**

| **Oligo name** | **Sequence** | **Purpose** |
|---|---|---|
| KTN1-seq-F | TAGATGGGCCCGAATCTAAAAT | KTN1 genomic fragment cloning to verify the mutation site |
| KTN1-seq-R | CTAAGTCTGAAGCCAAAAAACC | KTN1 genomic fragment cloning to verify the mutation site |
| KTN1g-F | CCATGATTACGAATTCTTCTTCTAGCTAGCTCTGGCAACC | KTN1::KTN1 cloning |
| KTN1g-R | TACGAACGAAAGCTCTGCAGGAAGCCAAAAAACCGAAGAAAATCATATAATCC | KTN1::KTN1 cloning |
| EN7-F | CCATGATTACGAATTCCTGCTCCATTAGTCCATATACACAGTTGAC | EN7::KTN1 and EN7::GFP-AGO1 cloning |
| EN7-R | CTAGAGGATCCCCGGGTACCTTTAAGATTCTGAGATTCACGAAGAAAAGC | EN7::KTN1 and EN7::GFP-AGO1 cloning |
| ACL5-F | CCATGATTACGAATTCTCGACGGCAAGCTATTCATTAGGCA | ACL5::KTN1 and ACL5::GFP-AGO1 cloning |
| ACL5-R | CTAGAGGATCCCCGGGTACCATCCAAGTTGAGGAGAAGATATAGAG | ACL5::KTN1 and ACL5::GFP-AGO1 cloning |
| GFP-F | ATCGGTACCATGATGGTGAGCAAGGGCGAGGA | EN7::GFP-AGO1, ACL5::GFP-AGO1 and AGO1::GFP-AGO1cloning |
| GFP-R | ATCGGATCCAGCTCCTCCTCCTCCCTTGTACAGCTCGTCCATGC | EN7::GFP-AGO1, ACL5::GFP-AGO1 and AGO1::GFP-AGO1cloning |
| AGO1-F | ATCCTCTAGAGTCGACATGGTGAGAAAGAGAAGAACGGATG | EN7::GFP-AGO1, ACL5::GFP-AGO1 and AGO1::GFP-AGO1cloning |
| AGO1-R | TACGAACGAAAGCTCTGCAGAGTTAAAGAAAGGATCAAAGTCTG | EN7::GFP-AGO1, ACL5::GFP-AGO1 and AGO1::GFP-AGO1cloning |
| AHP6-F | CCATGATTACGAATTCCACGGGGCGCAAAGAAGCATGACATAC | EN7::KTN1, AHP6:::KTN1, ACL5::KTN1 and SHR-KTN1 cloning |
| AHP6-R | CTAGAGGATCCCCGGGTACCCCACAACGGCACACCCGTCTTGTC | EN7::KTN1, AHP6:::KTN1, ACL5::KTN1 and SHR-KTN1 cloning |
| SHR-F | CCATGATTACGAATTCAGAAGCAGAGCGTGGGGTTTC | EN7::KTN1, AHP6:::KTN1, ACL5::KTN1 and SHR-KTN1 cloning |
| SHR-R | CTAGAGGATCCCCGGGTACCTTTTAATGAATAAGAAAATG | EN7::KTN1, AHP6:::KTN1, ACL5::KTN1 and SHR-KTN1 cloning |

| | | |
|---|---|---|
| KTN1-coding-F | ATCCTCTAGAGTCGACATGGTGGGAAGTAGTAATTCG | EN7::KTN1, AHP6:::KTN1, ACL5::KTN1 and SHR-KTN1 cloning |
| GR-F | CGGGGGACGAGCTCGGTACCATGGCTAGTGAAGCTCG AAAA | 35S::GR-GFP-AGO1 cloning |
| GR-R | TCGTATGGGTACATGGTACCAGCTCCTCCTCCTCCAGC TCCTCCTCCTCCGCTA GTAGATTTTTGATGAAACAG | 35S::GR-GFP-AGO1 cloning |
| GFP-R(GR) | ATCTCTAGACTTGTACAGCTCGTCCATGC | 35S::GR-GFP-AGO1 cloning |
| AGO1-F(GR) | AGGAGGAGCTTCTAGAGGAGGAGGAGGAGCTATGGT GAGAAAGAGAAGAACGG | 35S::GR-GFP-AGO1 cloning |
| AGO1-R(GR) | AGCTCTGCAGTCTAGAAGTTAAAGAAAGGATCAAAGT CTGT | 35S::GR-GFP-AGO1 cloning |
| AGO1p-F | CCATGATTACGAATTCTGACAGCCACCACATTCCTAA AG | AGO1::GFp-AGO1 cloning |
| AGO1p-R | CTAGAGGATCCCCGGGTACCGATGATTCCTGTGAAAA TAACAC | AGO1::GFp-AGO1 cloning |
| MIR165A-F | ATCCTCTAGAGTCGAC GTAAATCTACTCTTAAGAAGC | 35S::MIR165A cloning |
| MIR165A-R | TACGAACGAAAGCTCTGCAGCTACAACAAAAATTTGT GAATCTGC | 35S::MIR165A cloning |
| qSUL-F | GATCCAAAGATTGGTGGTGTTATG | real-time PCR |
| qSUL-R | AACTTGCTCTCCTTTCTCAACTCT | real-time PCR |
| qpri-MIR165A-F | CCATCATCACCATTCACCAACC | real-time PCR |
| qpri-MIR165A-R | CCAGACAACATTCCCCTCAACT | real-time PCR |
| qpri-MIR165B-F | TTTCTGTTGTGGGGAATGTTG | real-time PCR |
| qpri-MIR165B-R | CAGAGTGATTGAAGGCAATTAACAT | real-time PCR |
| qpri-MIR166A-F | TCTGGCTCGAGGACTCTGGC | real-time PCR |
| qpri-MIR166A-R | TGGAGTAAACAGGGAGCAACAAT | real-time PCR |
| qpri-MIR166B-F | TTTCTTTTGAGGGGACTGTTG | real-time PCR |
| qpri-MIR166B-R | ATTCGGATTGTGAGGGAGTG | real-time PCR |
| qpri-MIR166C-F | TGCGATTTAGTGTTGAGAGGATTG | real-time PCR |
| qpri-MIR166C-R | CGGTCGCAAGATAGAACAAATATGA | real-time PCR |
| qpri-MIR166D-F | GGTCATTTGCTCCTCTCTCCTATGT | real-time PCR |
| qpri-MIR166D-R | TCCTCTCAACCCTAAACCAAAGC | real-time PCR |
| qpri-MIR166E-F | GCTTTCTTGTCTCCTCCTCTTTCAG | real-time PCR |

| | | |
|---|---|---|
| qpri-MIR166E-R | GCCAGACAACATTCCCCTCAA | real-time PCR |
| qpri-MIR166F-F | CATGAGGCCGTAACTAAAAGAAAGTT | real-time PCR |
| qpri-MIR166F-R | TCTCGAGCCAGGCATCATTC | real-time PCR |
| qpri-MIR166G-F | TCTCTTTGCCCTGTCACATGCT | real-time PCR |
| qpri-MIR166G-R | TCCTCTAAACCCTAAATCGCTTCAC | real-time PCR |
| qATML1-F | TCCAGACGATAAGCAAAGAAAGG | real-time PCR |
| qATML1-R | CTTCAGTATCTGGTTCTCGTGCC | real-time PCR |
| qSUC2-F | GCACTAGCTTCCATATTTTCAACC | real-time PCR |
| qSUC2-R | GTCAACGCCAATACACCACTTAC | real-time PCR |
| qCAB3-F | GCCAAGCCAAAGGGTCCATCAGG | real-time PCR |
| qCAB3-R | TCCAGCGGTGTCCCATCCGTAGT | real-time PCR |
| N-UBQ5 | GGTGCTAAGAAGAGGAAGAAT | real-time PCR |
| C-UBQ5 | CTCCTTCTTTCTGGTAAACGT | real-time PCR |
| miR-SUL-as | AGGGATTTCCGTGACACTTAA | RNA gel blot |
| mir168-as | TTCCCGACCTGCACCAAGCGA | RNA gel blot |
| miR172-as | ATGCAGCATCATCAAGATTCT | RNA gel blot |
| miR167-as | TAGATCATGCTGGCAGCTTCA | RNA gel blot |
| miR166-as | GGGGAATGAAGCCTGGTCCGA | RNA gel blot |
| miR156-as | GTGCTCACTCTCTTCTGTCA | RNA gel blot |
| miR390-as | GGCGCTATCCCTCCTGAGCTT | RNA gel blot |
| miR169-as | TCGGCAAGTCATCCTTGGCTG | RNA gel blot |
| miR173-as | GTGATTTCTCTCTGCAAGCGAA | RNA gel blot |
| miR408-as | GCCAGGGAAGAGGCAGTGCAT | RNA gel blot |
| U6-as | AGGGGCCATGCTAATCTTCTCTG | RNA gel blot |
| tRNA-as | TCGAACTCTCGACCTCAGGAT | RNA gel blot |
